# Identifying SARS-CoV-2 Antiviral Compounds by Screening for Small Molecule Inhibitors of Nsp14/nsp10 Exoribonuclease

**DOI:** 10.1101/2021.04.07.438812

**Authors:** Berta Canal, Allison W. McClure, Joseph F. Curran, Mary Wu, Rachel Ulferts, Florian Weissmann, Jingkun Zeng, Agustina P. Bertolin, Jennifer C. Milligan, Souradeep Basu, Lucy S. Drury, Tom Deegan, Ryo Fujisawa, Emma L. Roberts, Clovis Basier, Karim Labib, Rupert Beale, Michael Howell, John F.X Diffley

**Author notes:** These authors contributed equally to this work (Berta Canal, Allison W. McClure, Joseph F. Curran).

## Abstract

SARS-CoV-2 is a coronavirus that emerged in 2019 and rapidly spread across the world causing a deadly pandemic with tremendous social and economic costs. Healthcare systems worldwide are under great pressure, and there is urgent need for effective antiviral treatments. The only currently approved antiviral treatment for COVID-19 is remdesivir, an inhibitor of viral genome replication. SARS-CoV-2 proliferation relies on the enzymatic activities of the non-structural proteins (nsp), which makes them interesting targets for the development of new antiviral treatments. With the aim to identify novel SARS-CoV-2 antivirals, we have purified the exoribonuclease/methyltransferase (nsp14) and its cofactor (nsp10) and developed biochemical assays compatible with high-throughput approaches to screen for exoribonuclease inhibitors. We have screened a library of over 5000 commercial compounds and identified patulin and aurintricarboxylic acid (ATA) as inhibitors of nsp14 exoribonuclease *in vitro*. We found that patulin and ATA inhibit replication of SARS-CoV-2 in a VERO E6 cell-culture model. These two new antiviral compounds will be valuable tools for further coronavirus research as well as potentially contributing to new therapeutic opportunities for COVID-19.

## Introduction

The severe acute respiratory syndrome coronavirus 2 (SARS-CoV-2) causes the human coronavirus disease 19 (COVID-19) [1, 2]. The large number of infections and the severe and long-term consequences of COVID-19 have greatly burdened healthcare systems worldwide [3–6]. Lockdowns and “stay-at-home” orders have been the only effective strategy to cut infections around the world, but they come with major societal and economic costs. Long term solutions to the health and economic crises will rely on our ability to properly monitor the evolution of the pandemic, on the effectiveness of novel vaccines, and on the development of new antiviral treatments to prevent the loss of further lives until we are able to control infections globally [7, 8].

Coronaviruses represent a threat to human health: in addition to SARS-CoV-2, two other coronaviruses, SARS-CoV-1 and MERS-CoV, have been responsible for severe human diseases this century [9]. Despite this, there is a lack of antiviral treatments for diseases caused by coronaviruses [10, 11]. Currently, the only antiviral agent approved by regulatory agencies for treatment of COVID-19 is remdesivir, a delayed chain-terminator nucleotide analogue that impairs SARS-CoV-2 viral replication [12]. However, the WHO-funded Solidarity trial failed to identify increased survival or reduced hospitalisation time in patients treated with remdesivir, casting doubt on its effectiveness in treating COVID-19 [13–16]. SARS-CoV-2 is a betacoronavirus of the order *Nidovirales* with a positive-sense RNA strand genome of approximately 30 Kb that encodes multiple open reading frames (ORF) [17, 18]. However, ORF1a and ORF1b represent two-thirds of the genome alone, and encode the 16 non-structural proteins of the virus (nsp1 - nsp16) [17]. The nsps include the 9 known viral enzymatic activities, which are highly conserved among coronaviruses and of special interest for the development of novel antiviral treatments, because of their important roles in viral replication and lifecycle [19].

Among them, nsp14 protein is a bifunctional enzyme with an N-terminal 3’ to 5’ exoribonuclease (ExoN) domain and a C-terminal S-adenosylmethionine (SAM)-dependent N7-methyltransferase (MTase) domain [20–22]. The ExoN domain belongs to the DEDD superfamily, which contains RNA and DNA exonucleases from all kingdoms of life, including the proofreading domains of some *E. coli* DNA polymerases [23–25]. While the MTase activity of nsp14 does not require a cofactor, the exonuclease activity of nsp14 is stimulated by the cofactor nsp10 [23, 26].

It has been suggested that the exonuclease activity of nsp14 may act as a proofreader to reduce the mutation rate of the virus [27–33]. Considering that most RNA viruses possess small genomes and have high mutation rates, proofreading activity could be important to maintain the integrity of the unusually large RNA genomes of coronaviruses including SARS-CoV-2. Consistent with the hypothesis that nsp14 plays a role in proofreading during replication, it associates with the viral RNA-dependent RNA polymerase (RdRp) complex formed by nsp12 and its cofactors nsp7 and nsp8 [34, 35]. Also, mutation of the ExoN domain of nsp14 in SARS-CoV-1 viruses resulted in viable viral progenies with high mutation rates [28, 33]. However, similar ExoN mutations led to inviable virus progenies in MERS-CoV and SARS-CoV-2, suggesting that the exonuclease activity of nsp14 could have other essential roles in viral replication in these viruses [36]. Furthermore, coronavirus nsp14 exonuclease activity has been proposed to reduce the host innate antiviral immune response by cleaving viral-associated double-stranded RNAs [37, 38] as well as to regulate viral genome recombination [39]. Thus, the exonuclease activity of nsp14 might be required for different aspects of viral genome replication and integrity and in other aspects of virus proliferation, making it an attractive target for the development of new antiviral treatments of COVID-19. Repurposing of previously characterised compounds for the treatment of novel diseases is a fast and efficient approach [40, 41]. Here, we have purified the SARS-CoV-2 nsp14/nsp10 exonuclease and screened a custom collection of previously characterised compounds with the aim to identify novel COVID-19 antivirals.

## Results

### Purified SARS-CoV-2 nsp14/10 functions as an exoribonuclease in vitro

To obtain sufficient quantities of active SARS-CoV-2 nsp14 exonuclease and nsp10 cofactor proteins for high-throughput screening, we tested a variety of protein tagging and expression strategies (Supplementary Table S1 and S2). All protein purifications involved an initial step based on an affinity tag followed by at least one other purification step (Supplementary Table S3). We first expressed and purified nsp14 and nsp10 individually from *Escherichia coli* (*E. coli or* ec) and baculovirus infected Sf9 insect cells (sf) (Figure 1A).

**Figure 1.**
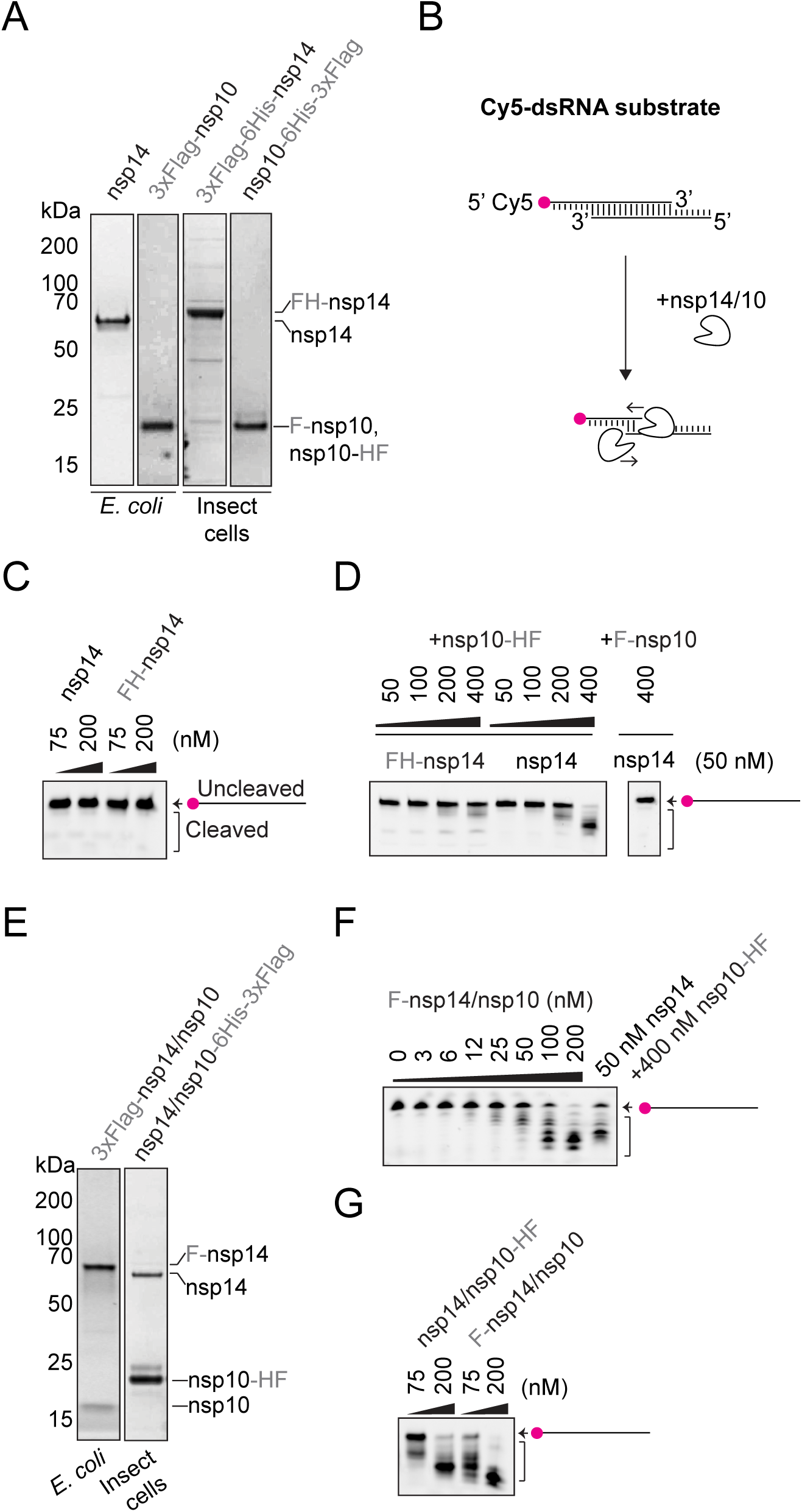
Purified nsp14/10 exoribonuclease assay. **A.** Coomassie-stained SDS-PAGEs of purified nsp14 (60 kDa) and nsp10 (15 kDa) proteins. **B.** Reaction scheme for gel-based exonuclease assay performed with the Cy5-substrate. Cleavage of the dsRNA substrate can be detected in TBE-urea polyacrylamide gels with the Cy5-substrate migrating faster as it is cleaved in the 3’ to 5’ direction. **C.** Nuclease reactions containing 75 nM or 200 nM nsp14 or Flag-His-nsp14 and 50 nM Cy5-substrate. Reactions performed at RT for 30 min visualised by TBE-urea polyacrylamide gels. **D.** As in C, first lane shows 400 nM Flag-nsp10 and remaining lanes show titration of nsp10-His-Flag over 50 nM nsp14. **E.** Purified co-expressed nsp14 and nsp10 complexes. **F.** As in C, titration of the co-expressed nsp14/10 complex and comparison to 50 nM nsp14 + 400 nM nsp10. **G.** As in C, reactions containing 75 nM or 200 nM of the co-expressed complexes nsp14/nsp10-His-Flag and Flag-nsp14/nsp10.

We tested nsp14 exoribonuclease activity using a double-strand RNA (dsRNA) substrate with 7-nucleotide overhangs on both the 5’ and 3’ ends, with one strand 5’ labelled with Cy5 (Cy5-dsRNA substrate) (Figure 1B). Reaction products were analysed on denaturing TBE-urea polyacrylamide gels, where generation of faster migrating products indicates enzymatic cleavage of the Cy5-dsRNA substrate. We did not observe cleavage of the Cy5-substrate in the presence of individual nsp14(ec) and 3xFlag-6His-nsp14(sf) proteins alone (Figure 1C). As shown previously [26], addition of increasing amounts of the cofactor nsp10 (nsp10-6His-3xFlag(sf)) promoted the nuclease activity of nsp14 and, to a lesser extent, of 3xFlag-6His-nsp14 (Figure 1D). In contrast to other reports [26], N-terminal tagged nsp10(ec) was unable to stimulate nsp14 activity at similar concentrations (Figure 1D).

We also purified co-expressed nsp14/10 complexes (Figure 1E). When the nsp14 subunit harboured the affinity tag for purification, the nsp10 subunit was less abundant and, reciprocally, when the nsp10 subunit harboured the affinity tag for purification, the nsp14 subunit was less abundant, suggesting that the complex formed by nsp14 and nsp10 is not stable stoichiometrically. Nevertheless, both nsp14/10 complexes co-expressed in *E.coli* or insect cells showed nuclease activity comparable to the single nsp14 + nsp10-6His-3xFlag proteins (Figure 1F and 1G).

### Quantitative fluorescence assay for nsp14/10 exoribonuclease activity

To study nsp14/10 exonuclease activity and perform a high-throughput screen, we developed a fluorescence-based assay using the intercalating Quant-it^TM^ RiboGreen RNA reagent (Figure 2A). We tested dilutions of RiboGreen against the Cy5-dsRNA substrate, and found a wide linear range of fluorescence using a 1/400 dilution of RiboGreen (Figure 2B and Supplementary Figure S1). RiboGreen fluorescence decayed after prolonged incubation in our reaction buffers (Figure 2C), so all subsequent experiments were analysed immediately after RiboGreen addition. To evaluate nsp14/10 exonuclease activity with the RiboGreen assay, we took the samples shown in Figure 1F and incubated them with RiboGreen. As predicted, RiboGreen fluorescence decreased with increasing amounts of nsp14/10 consistent with exoribonuclease activity (Figure 2D).

**Figure 2.**
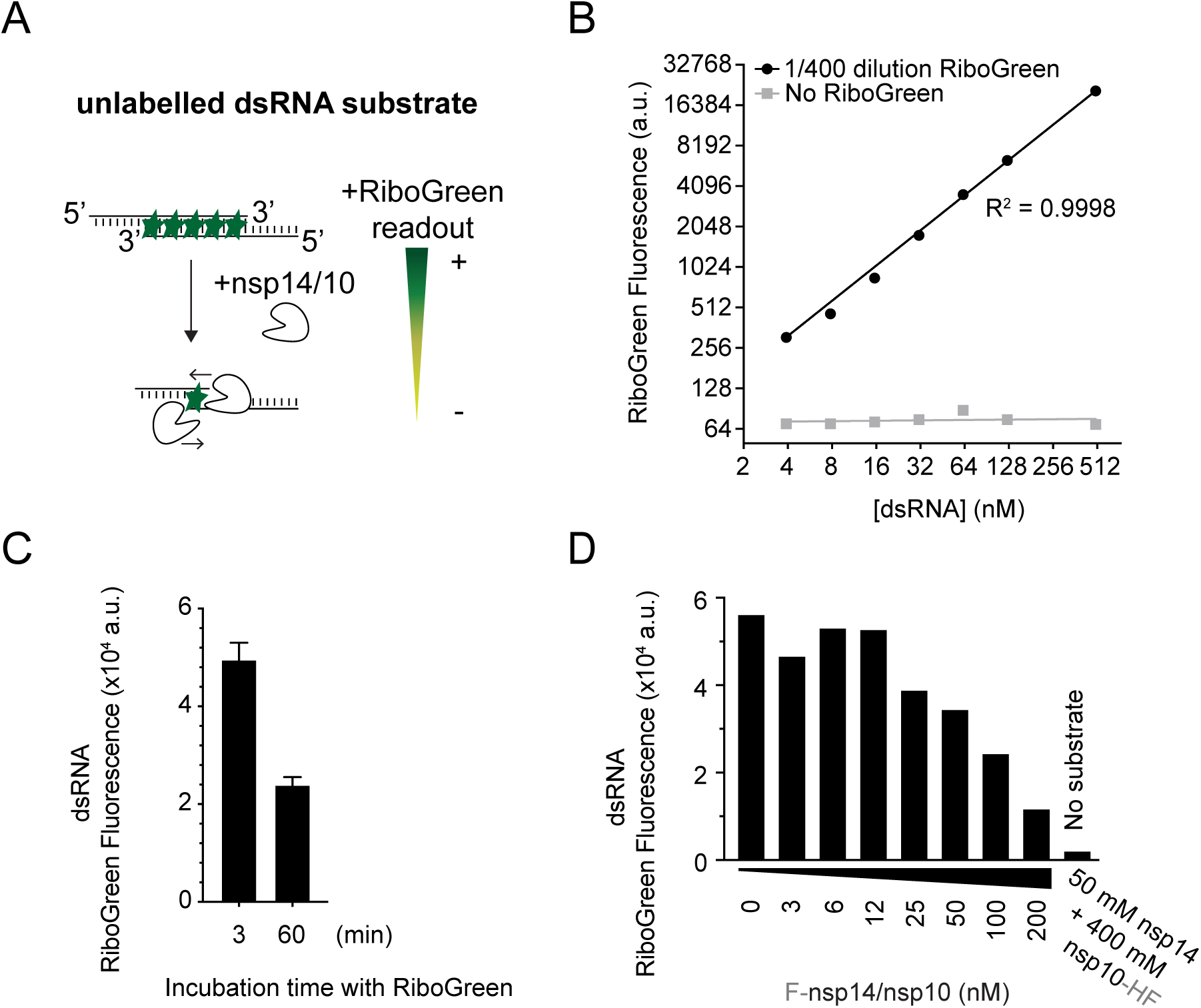
RiboGreen assay for nsp14/10 exoribonuclease activity. **A.** Reaction scheme for exonuclease activity detected by RiboGreen fluorescence. **B.** Titration of the Cy5-dsRNA substrate incubated with RiboGreen and detected by microplate reader. **C.** Cy5-dsRNA substrate was incubated with RiboGreen for 3 min or 60 min prior to fluorescence detection. Error bars represent standard deviation from the mean. **D.** Exonuclease reactions from Figure 1F. 30 min reactions with titration of the co-expressed nsp14/10 complex and 50 nM of Cy5-substrate and comparison to 50 nM nsp14 + 400 nM nsp10, followed by RiboGreen incubation.

### nsp10-14 fusion functions as an efficient exoribonuclease

The untagged subunit in the nsp14/10 complex preparations tended to be substoichiometric (Figure 1E), and relatively high amounts of nsp10 were required to stimulate nsp14 (Figure 1D), suggesting that the nsp14/10 complex might not be stable. We therefore wondered if direct fusion of nsp10 to nsp14 would generate a more active enzyme. A similar strategy was successful with the SARS-CoV-1 proteins nsp12/7/8, which purify as a more active polymerase complex when nsp7 and nsp8 are fused together with a short linker [34]. To test this, we expressed nsp14-linker-nsp10 (nsp14-10) and nsp10-linker-nsp14 (nsp10-14) fusion proteins in *E. coli* (Figure 3A and 3B). The nsp10-14 fusion cleaved the Cy5-dsRNA substrate in both the gel and RiboGreen assays (Figure 3C). Titration of nsp10-14 fusion protein showed high exonuclease activity with as little as 1-3 nM of protein compared to about 200 nM needed for the nsp14/10 co-expressed complex (Figure 3D). The nsp14-10 fusion protein also showed high exonuclease activity (Figure 3D). To avoid any interference between Cy5 and RiboGreen, we then titrated the nsp10-14 fusion in the presence of an unlabelled version of the Cy5-dsRNA substrate (Figure 2A and Supplementary Table S4), which resulted in exonuclease activity (Figure 3E) comparable to that observed with the Cy5 substrate (Figure 3D). Nsp14 is also an N7-guanine cap methyltransferase and the nsp10-14 fusion retains this methyltransferase activity (Supplementary Figure S2).

**Figure 3.**
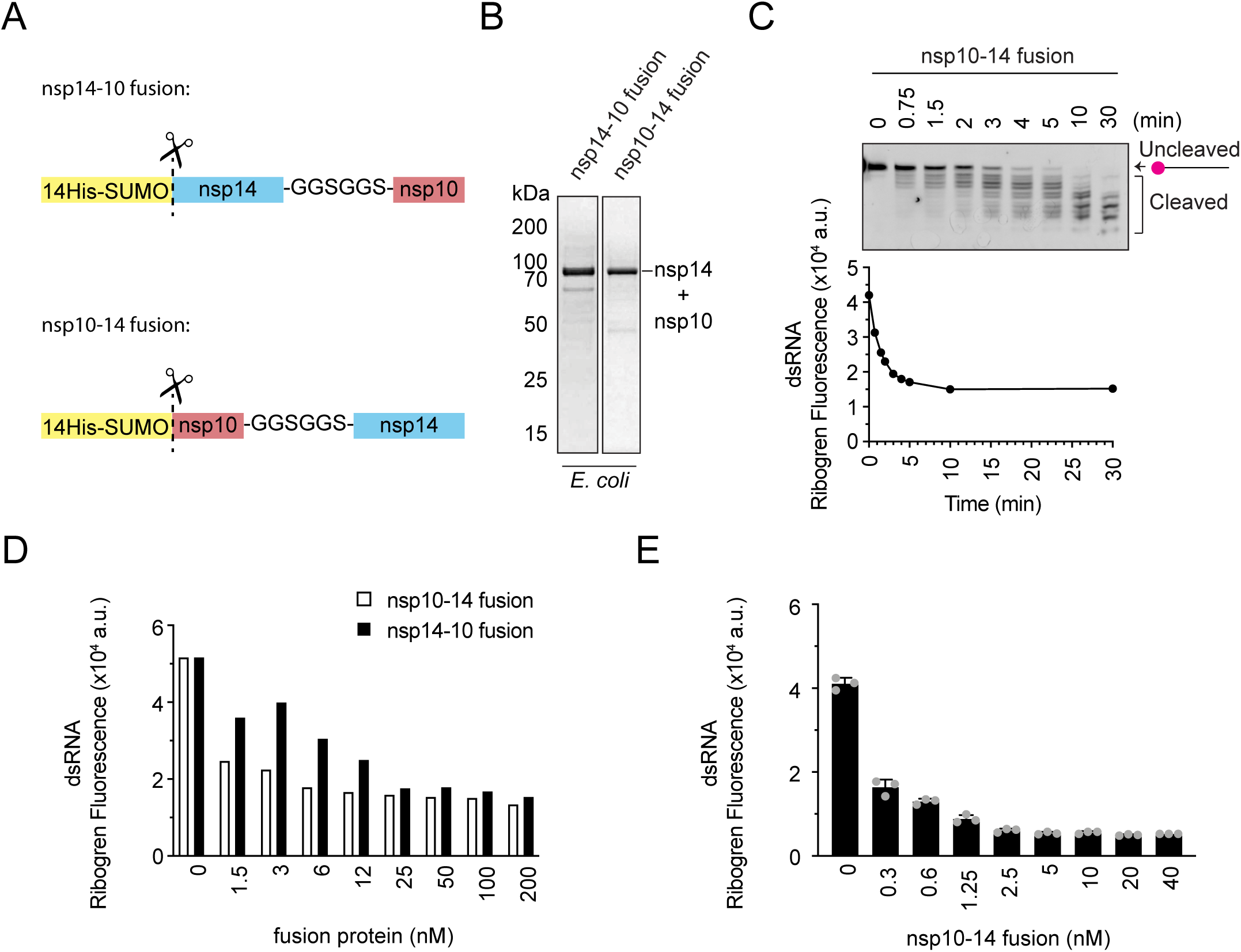
The nsp14-10 fusion protein is an efficient exonuclease. **A.** Schematic of nsp14-10 and nsp10-14 fusion proteins. **B.** Coomassie-stained SDS-PAGEs of purified nsp14-10 and nsp10-14 fusion proteins (75 kDa). Connecting linker is GGSGGS. **C.** Time course of exonuclease reaction containing 5 nM nsp10-14 fusion and 50 nM Cy5-substrate. Reactions were split in half and separated on TBE-urea polyacrylamide gels or incubated with RiboGreen. **D.** Exonuclease reactions containing a titration of nsp10-14 and nsp14-10 fusion proteins and 50 nM Cy5-substrate performed for 30 minutes. **E.** Titration of nsp10-14 fusion in exonuclease assay with 50 nM unlabelled dsRNA substrate after 30 min detected by RiboGreen. Error bars represent standard deviation from the mean.

### Screens for SARS-CoV-2 nsp14/10 inhibitors

The quantitative nature of the RiboGreen assay enabled us to determine the Michaelis-Menten constant (K_M_) of the nsp10-14 fusion protein for the unlabelled dsRNA substrate to be 66 nM (Supplementary Figure S3A and S3B). We then carried out a high-throughput screen to identify nsp14/10 exonuclease inhibitors from a custom library of over 5000 commercial compounds. The screen had a robust signal to noise ratio between positive and negative controls, and a mean Z’ factor of 0.72, and we detected many compounds apparently able to reduce nsp14/10 exonuclease activity at both concentrations screened (Supplementary Figure S3C). In the RiboGreen assay, fluorescence is highest at time zero of the exonuclease reaction, and cleavage of the substrate leads to reduced fluorescence signals (Figure 2D or 3D as examples). Thus, inhibition of exonuclease activity is expected to decrease the cleavage-dependent reduction in RiboGreen fluorescence. Based on this, to identify putative inhibitors of nsp14/10 exonuclease, we selected compounds that led to higher final fluorescence signals than control reactions. However, when we were validating these compounds, we found that many were auto-fluorescent in the wavelength range of RiboGreen fluorescence (Supplementary Figure S3D). Because of this, we could not feasibly distinguish true hit compounds from auto-fluorescent false hits using this approach.

We decided to perform a second fluorescence-based high-throughput screen against the same library but using a different assay which utilises a different wavelength. For the new assay, we used an RNA duplex substrate with a 20-nucleotide 5’ dU overhang, conjugated to a Cy3-quencher pair (Figure 4A and Supplementary Table S4). Using this substrate, nucleolytic cleavage by nsp10-14 fusion protein released the Cy3 fluorophore strand from the quencher strand and produced fluorescent signal (Figure 4A and 4B). When a DNA version of this substrate was used, only background levels of fluorescence were detected (Figure 4B). This indicated that, as expected, nsp10-14 is unable to cleave DNA substrates and that the fluorescence in reactions with the Cy3-dsRNA substrate is not due to unspecific factors such as temperature-dependent unwinding. When using this substrate, we were able to obtain kinetic data of the enzyme compared to the end-point information obtained when using our previous RiboGreen approach. We, thus, monitored fluorescence over time and titrated the substrate, which gave an estimated K_M_ of 40 nM (Figure 4C and 4D). The estimated K_M_ was similar to our previous estimation using the RiboGreen assay (66 nM) (Supplementary Figure S3A and S3B), and therefore we decided to perform this second screen using 50 nM of substrate, as we did in the first screen. At 50 nM of substrate, we assessed nuclease activity over a range of enzyme concentrations and chose to do the reactions in the screen for 10 minutes with 0.5 nM of nsp10-14 fusion protein (Figure 4E).

**Figure 4.**
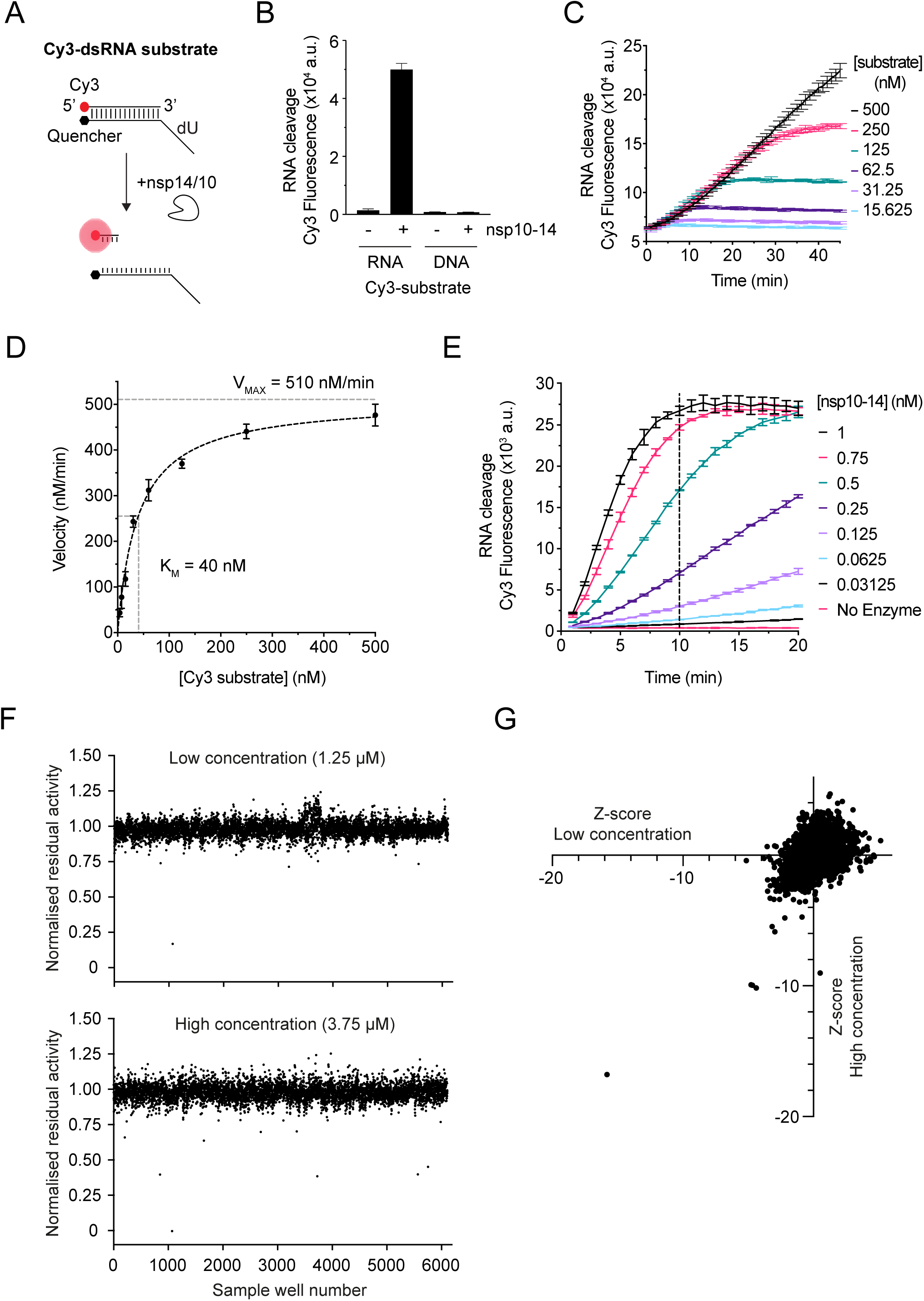
High-throughput screen to identify SARS-CoV-2 nsp14/10 exoribonuclease inhibitors. **A.** Reaction scheme for the kinetic nuclease assay performed with the Cy3-dsRNA substrate. Cleavage of the dsRNA substrate by nsp14/nsp10 nuclease can be detected by the appearance of Cy3 fluorescence. **B.** Exonuclease reactions containing 5 nM nsp10-14 fusion protein and 180 nM Cy3-dsRNA substrate or 180 nM Cy3-dsDNA substrate and detected after 1 h. **C.** Cy3-dsRNA substrate titration (15 - 500 nM) performed in the presence of 0.5 nM nsp10-14 with fluorescence monitored over time. **D.** Non-linear fit to Michaelis-Menten equation of slopes from C. **E.** Titration of nsp10-14 with 50 nM Cy3-dsRNA substrate. **F.** Normalised residual activity of screen sample wells in low and high concentrations. **G**. Z-scores of samples plotted as low versus high concentration. In all panels, error bars represent standard deviation from the mean.

As performed in the first screen, the drug library was screened at 1.25 µM and 3.75 µM. First, nsp10-14 was incubated with the compounds for 10 min prior to the addition of the substrate, then fluorescent measurements were taken every minute for 10 min. The activity of nsp10-14 was defined as the slope of the reaction over the linear range. Again, the screen had a robust signal to noise ratio between positive and negative controls and a mean Z’ factor of 0.86. The activity of nsp10-14 was normalised to the control wells without compounds (see Experimental Procedures). Both compound concentrations produced hits with residual activities less than 1 (Figure 4F). We considered compounds that reduced nsp10-14 activity below 80% in both concentrations, showed a Z score less than −5 (Figure 4G), and also showed some reduction in the first screen with RiboGreen detection. We also considered compounds that showed stronger inhibition in the higher dose in both screens, even if the absolute inhibition was not as strong as our initial cut-offs. We discounted some compounds for having a high aggregation index (>3 or known aggregators) [42] and for appearing in multiple screens (see accompanying manuscripts).

### Validation of hits

Taking the results from both screens into consideration, we tested 12 compounds in validation experiments (Supplementary Table S5). We returned to the gel-based assay with the Cy5-dsRNA substrate for validation. The nsp10-14 fusion protein was incubated with the hit compounds for 10 minutes prior to substrate addition as in the screens, and the reactions were stopped at 5 min instead of the usual 30 min reactions, to allow for the visualization of both the full-length RNA strand and the cleaved products. Two compounds inhibited nsp10-14 exonuclease activity in a concentration-dependent manner at both 5 µM and 25 µM: patulin and aurintricarboxylic acid (ATA) (Figure 5A). We also tested whether patulin inhibited the co-purified complex nsp14/10 in addition to nsp10-14 fusion protein. Indeed, patulin inhibited both fusion and complex when in ∼ 5000 fold excess of enzyme (Figure 5B, 5C lanes 2 - 5, and Supplementary Figure S4A).

**Figure 5.**
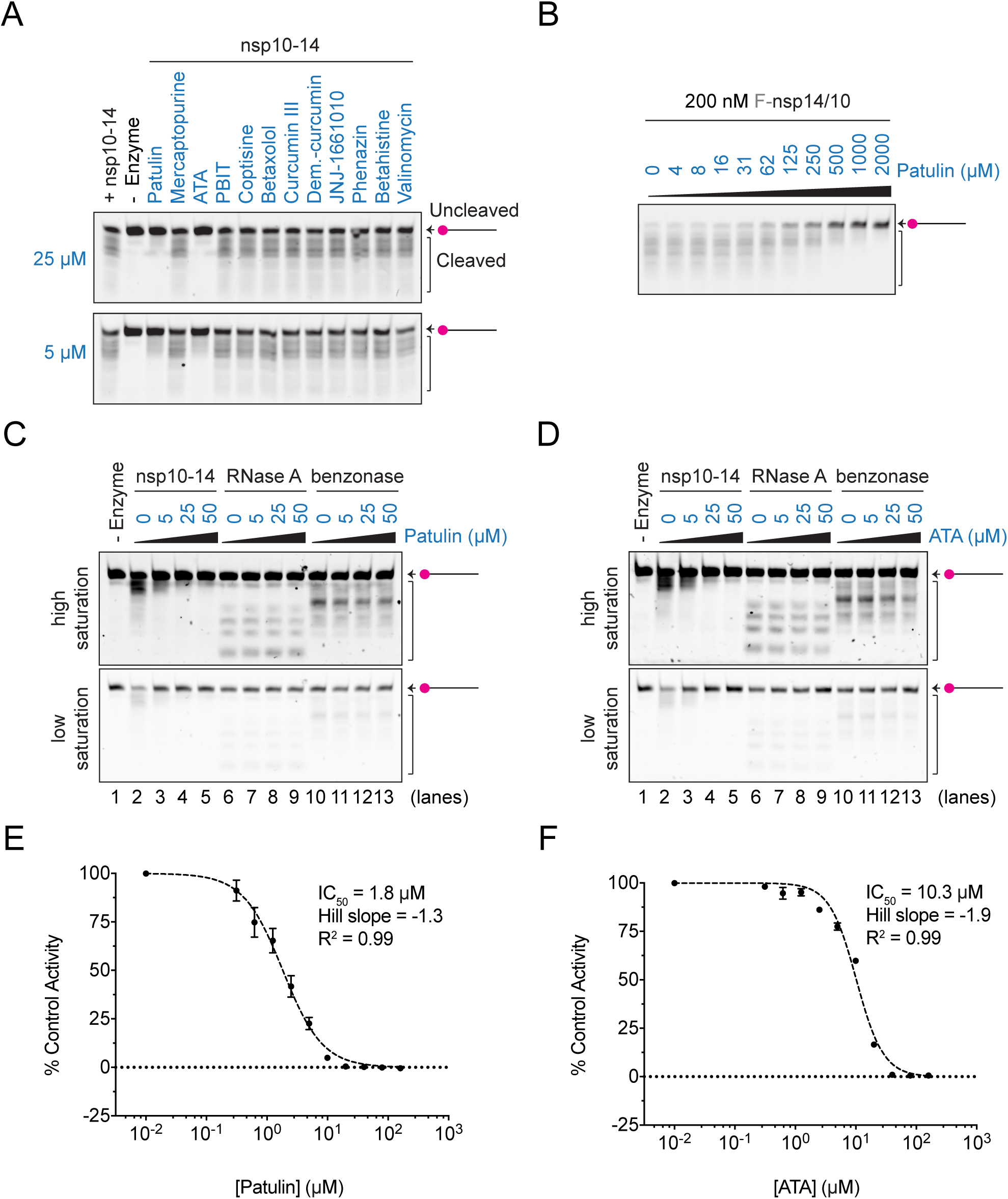
Patulin and aurintricarboxylic acid (ATA) inhibit nsp14/10 nuclease in vitro. **A.** Nuclease reactions containing 1 nM nsp10-14 fusion protein and 50 nM of Cy5-dsRNA substrate alone or in the presence of 5 µM or 25 µM of each of the 12 top hits (in blue) selected from both HTS screens. Reactions were performed for 5 minutes at RT and visualised by TBE-urea polyacrylamide gels. **B.** 200 nM co-expressed nsp14/10 complex was pre-incubated with the specified concentration of patulin, and nuclease reactions were performed in the presence of 50 nM Cy5-dsRNA substrate for 5 min and visualised by TBE-urea polyacrylamide gels. **C, D.** Nuclease reactions containing 1 nM nsp10-14 fusion protein, 0.06 ng/ul RNase A (Qiagen) and 0.5 mU/ul benzonase (Sigma) nucleases and 50 nM Cy5-dsRNA substrate in the presence of 0, 5, 25 or 50 µM patulin (C) or ATA (D). High and low image saturations are shown. **E, F.** Dose-response curves and IC_50_ values of patulin (E) and ATA (F). IC_50_ values were calculated as described in Experimental Procedures. Nuclease activities (slope) for each patulin and ATA concentration were obtained from triplicate kinetic reactions (20 min) in the presence of 0.5 nM nsp10-14 fusion and 50 nM Cy3-substrate (Supplementary Figure S4E, S5F). Patulin and ATA were pre-incubated with nsp10-14 for 10 minutes prior to the addition of the substrate. Graphs represent % of control activity (slope) vs. the log_10_ of the concentration of patulin or ATA. In E and F, error bars represent standard deviation from the mean.

To determine how specific patulin and ATA are to nsp10-14, we tested whether they inhibited other nucleases such as RNase A and benzonase. We titrated RNase A and benzonase to select enzyme concentrations that allowed us to visualise both full-length RNA and cleaved products in gel-based assays after 5 min (Supplementary Figure S4B). Patulin did not inhibit RNase A or benzonase up to 50 µM, suggesting that it is not a general nuclease inhibitor (Figure 5C). Despite the fact that ATA has been described as a general inhibitor of nucleases and other DNA binding enzymes [43], we observed only minor inhibition of RNase A and benzonase at 50 µM ATA (Figure 5D). In contrast, we observed inhibition of nsp10-14 from 5 µM ATA, indicating that it inhibits nsp10-14 at lower concentrations than it inhibits RNase A and benzonase (Figure 5D).

To characterise the inhibition kinetics of patulin and ATA (chemical structures in Supplementary Figure S4C and S4D) on nsp14/10 exonuclease, we determined the half maximal inhibitory concentration (IC_50_) of these compounds. To calculate the IC_50s_, we incubated increasing concentrations of patulin or ATA with nsp10-14 protein following the scheme of the screen and calculated the exonuclease activity of nsp10-14 at each drug concentration in the presence of the Cy3-dsRNA substrate (see Experimental Procedures). Patulin showed an IC_50_ of 1.8 µM (95% CI 1.6 - 2.1 µM) and ATA showed an IC_50_ of 10.3 µM (95% CI 8.6 - 12.1 µM) (Figure 5E and 5F and Supplementary Figures S4E and S4F). Importantly, in all the concentrations assayed, patulin and ATA do not quench Cy3 fluorescence (Supplementary Figure S4G and S4H).

### Effects of patulin and aurintricarboxylic acid on viral growth

To explore if patulin and ATA could serve as antiviral drugs against SARS-CoV-2, we performed infectivity assays using VERO E6 cells in the presence of increasing concentrations of patulin and ATA. SARS-CoV-2 nsp14 exonuclease activity has been recently proposed to be essential for viral proliferation, and thus we expected that if patulin and ATA were able to inhibit nsp14 exonuclease also in cells, we would observe reduced viral infectivity [36]. Virus in VERO E6 cells 22 hours post-infection (MOI 0.5 PFU/cell) was detected by immunofluorescence with an antibody raised against the SARS-CoV-2 nucleocapsid (N) protein (see Experimental Procedures). Viral proliferation was reduced in the presence of patulin at ∼ 5-10 µM (Figure 6A and 6B). Above 10 µM, patulin decreased cell viability of cultured VERO E6 (Figure 6A and 6B). Similarly, we observed reduced viral proliferation in the presence of ATA in a dose-dependent manner from ∼ 3-100 µM (Figure 6C). At the concentrations tested, ATA did not affect cell viability (Figure 6C).

**Figure 6.**
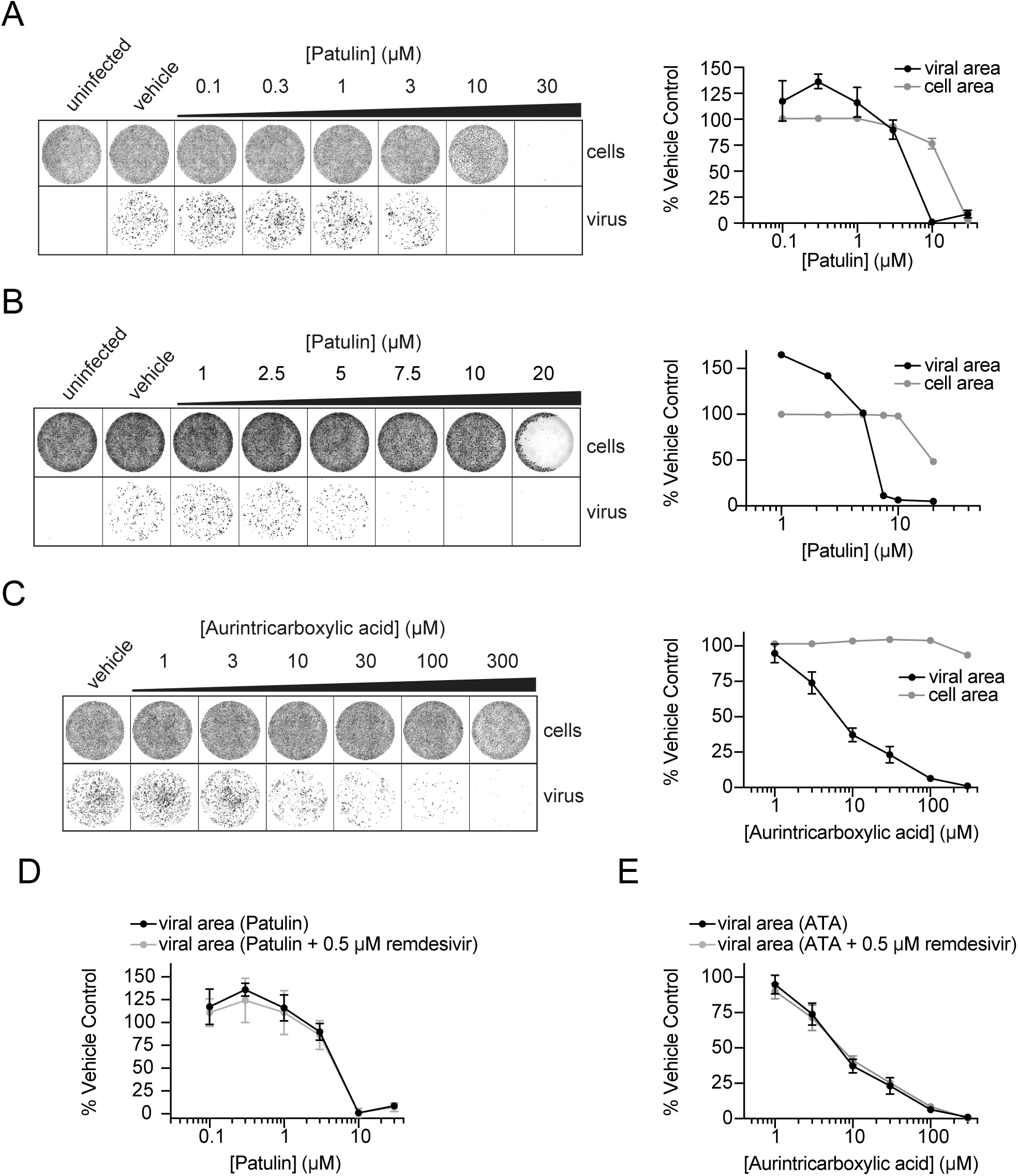
Patulin and aurintricarboxylic acid (ATA) inhibit SARS-CoV-2 viral growth in VERO E6 cells. **A.** After VERO E6 cell seeding, media was replaced with fresh media containing indicated concentrations of patulin followed by infection with SARS-CoV-2. 22 h post-infection, wells were stained with DR (cells) and AlexaFluor-conjugated antibodies against the SARS-CoV-2 N protein (virus). Quantification is shown at right (mean +/-SD). Quantification of viral area is normalised to cell area. **B.** Experiment performed as in A with lower dose titration of patulin. **C.** Experiment as in A, with titration of aurintricarboxylic acid (ATA). **D.** Experiment as in A, with combined treatment of 0.5 µM remdesivir. Note the black line indicating patulin only treatment is the same as in A. **E.** Experiment as in C, with combined treatment of 0.5 µM remdesivir. Note the black line indicating ATA only treatment is the same as in C.

Combining different antivirals is effective in treating viral infections and reducing treatment resistance [44]. To this end, we performed the SARS-CoV-2 infectivity assays in the presence of patulin or ATA in combination with the SARS-CoV-2 antiviral remdesivir. When combined, the dose-response curves of patulin and ATA were not shifted in response to a low dose (0.5 µM) of remdesivir, indicating that, under the conditions tested, these drugs do not synergise (Figure 6D and 6E). We also performed the reciprocal experiment testing whether patulin would alter the remdesivir dose-response curve, but we did not observe any shift in the remdesivir curve in the presence of 3 µM or 5 µM patulin (Supplementary Figure S5).

## Discussion

Previously, nsp14/nsp10 exonuclease activity had only been studied by monitoring cleavage of RNA substrates in gel-based assays using an excess of enzyme over substrate concentration [26, 36, 45, 46]. In this study, we have purified a fusion protein of nsp14 and nsp10 that shows ∼ 100 fold higher exoribonuclease activity than the co-expressed or single nsp14 and nsp10 proteins. Further, we developed fluorescence-based assays for nsp14/10 exoribonuclease activity in solution, which are also scalable for high-throughput screens. We used these assays to screen for nsp14/10 exoribonuclease inhibitors among a custom chemical library of over 5000 commercial compounds and identified patulin and aurintricarboxylic acid (ATA) as inhibitors of SARS-CoV-2 nsp14/10 exonuclease.

The exoribonuclease (ExoN) domain of nsp14 is highly conserved among the nidoviruses that possess large genomes (>20 Kb), however, the function of ExoN for viral proliferation is still unclear. In addition to its proposed role as proofreader, nsp14/10 is thought to promote host innate immune evasion by cleaving dsRNA molecules and participate in viral genome recombination [37, 39, 47, 48]. ExoN is essential in SARS-CoV-2 [36], so patulin and aurintricarboxylic acid (ATA), the new inhibitors of nsp14/10 exonuclease activity identified here, represent new tools to acutely inhibit ExoN activity during viral growth, which should be valuable for understanding the role of nsp14/10 ExoN during the viral life cycle. Also, considering the role of nsp14/10 in immune suppression, we speculate that even mild inhibition of nsp14/10 ExoN by patulin and ATA could lead to accumulation of highly immunogenic dsRNA molecules and promote strong immune antiviral responses.

A proofreader activity of nsp14 could reduce the effectiveness of nucleotide analogue chain terminators such as remdesivir by preventing their incorporation or removing them [49]. Supporting this, ExoN mutants of murine hepatitis virus (MHV), a type of coronavirus, were shown to have 4.5-fold higher sensitivity to remdesivir treatment [12]. However, our results in VERO E6 cells did not show any synergy of patulin or ATA with remdesivir (Figure 6C and 6D). Remdesivir acts as a delayed chain terminator where RNA synthesis stops about 3 nucleotides after its incorporation and perhaps nsp14/10 does not have access to remove it. It would be interesting for future studies to examine whether patulin and ATA synergise with immediate chain terminators and to further explore how nsp14/10 might act as a proofreader.

ATA has been used as a positive control drug for nuclease inhibition due to its known roles as a general nuclease and DNA binding inhibitor [43, 50]. Indeed, ATA has recently been used as a control to inhibit nsp14/nsp10 exoribonuclease using a biochemical gel-based assay [51]. However, our results show that ATA inhibits nsp14/nsp10 exoribonuclease activity at much lower concentrations than the concentration needed to inhibit other nucleases such as RNase A and benzonase (Figure 5D), suggesting that ATA has specific activity towards SARS-CoV-2 nsp14/nsp10 compared to other nucleases. We cannot rule out the possibility that some of the antiviral activity of ATA is due to inhibition of other nucleases besides nsp14/10. ATA has been shown to inhibit replication of several other viruses including SARS-CoV-1 and Zika [52, 53], and studies using cell culture and animal models have shown limited toxicity of ATA [53–55]. Further studies will be needed to understand the putative use of ATA for the inhibition of SARS-CoV-2 and treatment of COVID-19.

Patulin is a lactone mycotoxin produced by several species of *Penicillium* and *Aspergillus*, commonly found in foodstuffs such as rotting apples [56]. Patulin has antimicrobial effects and it has been shown to have cytotoxic, apoptotic and genotoxic effects in mammalian cells, including cancer cells [56–59]. While patulin does show some cellular toxicity in our assays, structural analogues of patulin were shown to have reduced toxicity compared to patulin in cell culture models [60], providing an opportunity to study alternative patulin-like molecules for the development of novel antiviral agents against coronaviruses. Also, considering that even mild inhibition of nsp14/10 exonuclease could have an effect on host immune responses, low and transient doses of patulin could also be considered. After some initially promising results in the early 20^th^ century, patulin was tested in a clinical trial for its ability to treat the common cold but found to be ineffective [61]. Since the time of this MRC trial, it has been shown that many viruses in addition to coronaviruses can cause the common cold, so it is likely that if patulin was effective only in treating coronavirus-based colds, the trial would not have uncovered it. It will be interesting for future studies to test patulin in the treatment of both COVID-19 and other coronavirus diseases.

## Materials and Methods

### Expression constructs and protein expression in E.coli

The sequences of SARS-CoV-2 nsp14 and nsp10 (NCBI reference sequence NC_045512.2) were codon optimised for expression in E. coli using the GeneArt Gene Synthesis software (ThermoFisher scientific) and genes were ordered from GeneWiz (codon optimised DNA sequences in supplementary information). The HiFi DNA Assembly system (NEB) was used to build the plasmids to express individual, complex and fusion proteins (Supplementary Table S1 and S3). <u>Individual nsp10</u> was expressed from plasmid SARS-CoV-2 3xFlag-nsp5CS-nsp10 (Addgene ID 169157) containing the N-terminal 3xFlag tag (MDYKDHDGDYKDHDIDYKDDDDK). Untagged <u>individual nsp14</u> (nsp14) was expressed from plasmid DU70487 that contains non-codon optimised nsp14 gene cloned into plasmid pK27SUMO (gene synthesis at MRC PPU available from https://mrcppu-covid.bio). To co-express <u>nsp14/10 complex</u>, we co-transformed plasmids SARS-CoV-2 nsp10 (Addgene ID 169158) and SARS-CoV-2 3xFlag-nsp5CS-nsp14 (Addgene ID 169159). To express nsp10-14 and nsp14-10 <u>fusion proteins</u>, nsp10 and nsp14 genes from plasmids 169157 and 169159 were cloned in frame with the N-terminal 14His-K27SUMO tag of plasmid pK27Sumo [62], including a GGSGGS linker between both nsp proteins (Figure 3A). For expression of the nsp10-14 fusion protein, T7 Express LysY competent cells from NEB (C3010I) were transformed with plasmid pEcoli-Cov_42 (Supplementary Table S2) and grown overnight (ON) at 37°C in LB media supplemented with 50 μg/ml kanamycin. Saturated cultures were diluted in 10 L to an OD_595_ of 0.1 and grown at 37°C until OD_595_ of 0.4. Cultures were then cooled down to 16°C on ice, after which protein expression was induced ON at 16°C by adding 0.4 mM IPTG. Cells were harvested by centrifugation, washed once in 1x TBS, and the pellet was frozen in liquid nitrogen. All steps after this point were carried out at 4°C. Pellets were resuspended to 30 mL with lysis buffer (50 mM Tris-HCl pH 7.5, 0.05 % NP-40, 10 % glycerol, 500 mM NaCl, 1 mM DTT and 30 mM imidazole) + cOmplete™ EDTA-free Protease Inhibitor Cocktail (Sigma). Cells were lysed for 30 min on ice by addition of 0.33 mg/ml lysozyme followed by sonication (5 sec on/5 sec off at 35% for 3 min), and cell debris was cleared by centrifugation at 20,000 rpm for 30 min.

### Expression constructs and protein expression in baculovirus-infected insect cells

Plasmids SARS-CoV-2 nsp10-His-3xFlag (Addgene ID 169162), SARS-CoV-2 3xFlag-His-nsp5CS-nsp14 (Addgene ID 169163) and SARS-CoV-2 nsp14/nsp10-His-3xFlag (Addgene ID 169164) were expressed in baculovirus-infected Sf9 insect cells. The coding sequences of SARS-CoV-2 nsp10 and nsp14 (NCBI reference sequence NC_045512.2) were codon-optimised for *S. frugiperda* ordered from GeneArt (Thermo Fisher Scientific) (codon optimised DNA sequences can be found in supplementary information). Baculoviral expression vectors were generated using the biGBac vector system [63]. Nsp10 was subcloned into a modified pBIG1b vector containing a pLIB-derived polyhedrin expression cassette to contain a C-terminal 6His-3xFlag tag (sequence: nsp10-GGSHHHHHHGSDYKDHDGDYKDHDIDYKDDDDK). Nsp14 was subcloned into a modified pBIG1a vector containing a pLIB-derived polyhedrin expression cassette to either contain an N-terminal 3xFlag-6His tag (sequence: MDYKDHDGDYKDHDIDYKDDDDKGSHHHHHHSAVLQ-nsp14) or no tag to generate SARS-CoV-2 nsp14/nsp10-His-3xFlag (Addgene ID 169164). Baculoviruses were generated and amplified in Sf9 insect cells (Thermo Fisher Scientific) using the EMBacY baculoviral genome [64]. For protein expression, Sf9 insect cells were infected with baculovirus and collected 48 h after infection, flash-frozen and stored at −70 °C. Pellets were lysed with a Dounce homogeniser, 3 x 10 strokes on ice and cell debris cleared by centrifugation at 20,000 rpm for 30 min.

### Protein purification

For the purification of the nsp10-14 fusion protein, 1 ml slurry of HisPur^TM^ Ni-NTA resin equilibrated in lysis buffer was added to the cleared lysate and rotated for 2 h. Resin was collected by centrifugation at 1000 x g for 2 min, washed with 100 ml lysis buffer and incubated for 20 min in 10 ml elution buffer (lysis buffer with 400 mM imidazole). Eluate was dialysed ON against 1.5 L of dialysis buffer (25 mM Tris-HCl pH 7.5, 0.02 % NP-40, 10 % glycerol, 100 mM NaCl, 1 mM DTT and 30 mM imidazole) + 20 μg/ml Ulp1-cat-6His, and passed over 0.5 ml HisPur^TM^ Ni-NTA resin pre-equilibrated in dialysis buffer to remove Ulp1. The flowthrough was applied to a MonoQ (5/50 GL, GE healthcare) equilibrated in Buffer 100 (25 mM Tris-HCl pH 7.5, 0.02 % NP-40, 10 % glycerol, 1 mM DTT and 100 mM NaCl), washed with 10 CV Buffer 100 and eluted over a 20 CV linear gradient to Buffer 1000 (25 mM Tris-HCl pH 7.5, 0.02 % NP-40, 10 % glycerol, 1 mM DTT and 1000 mM NaCl). The nsp14-10 and nsp10-14 fusion proteins eluted at ∼ 400 mM NaCl. Peak fractions were pooled, concentrated to ∼ 0.4 ml with a 30KDa Amicon Ultra Centrifugal filter and separated on a Superdex 200 Increase 10/300 GL column equilibrated in Buffer 150 (25 mM Tris-HCl pH 7.5, 0.02 % NP-40, 10 % glycerol, 1 mM DTT, 4 mM MgCl2 and 150 mM NaCl). Peak fractions were pooled, aliquoted and frozen in liquid nitrogen to be stored at −80°C. Protein concentration was determined by Bradford and by comparing to a BSA standard curve on a Coomassie stained SDS-PAGE gel. The nsp10-14 fusion protein concentration was at 1.1 mg/ml (yield ∼ 0.4 mg protein / L cell culture).

Modifications of this protocol were used to purify the other recombinant proteins (Supplementary Table S3). Purifications using 3xFlag tag were performed similarly with the following modifications: 1 ml slurry of ANTI-FLAG M2 Affinity gel (Sigma) equilibrated in flag lysis buffer (50 mM Tris-HCl pH 7.5, 0.05 % NP-40, 10 % glycerol, 500 mM NaCl, 1 mM DTT) was added to the cleared lysate and rotated for 2 h. Resin was collected by centrifugation at 1000 x g for 2 min, washed with 100 ml flag lysis buffer and incubated for 30 min in 10 ml flag elution buffer (50 mM Tris-HCl pH 7.5, 0.05 % NP-40, 10 % glycerol, 300 mM NaCl, 1 mM DTT) + 0.5 mg/ml 3xFLAG peptide. Eluate was diluted 3-fold with No salt buffer (50 mM Tris-HCl pH 7.5, 0.05 % NP-40, 10 % glycerol, 1 mM DTT) to be applied to a MonoQ or concentrated to ∼ 0.4 ml with a 30KDa Amicon filter for gel filtration.

### Expression and purification of Ulp1-cat-6His protease

T7 Express *LysY* competent cells transformed with plasmid pFGET19-Ulp1 (Addgene 64697) were grown to OD_595_ of 0.8 at 37°C in 2 L LB media supplemented with 50 μg/ml kanamycin (ON culture at 37°C without shaking). Expression was induced for 4 h at 37°C with shaking by the addition of 1 mM IPTG. Pellet was resuspended in lysis buffer (50 mM Tris-HCl pH 8, 0.02% NP40, 10% Glycerol, 500 mM NaCl, 5 mM MgOAc, 1 mM DTT) + 20 mM imidazole + protease inhibitors (10 μg/ml Pepstatin -P4265 Sigma-, 10 μg/ml Leupeptin −108975 Merck-, 1 mM AEBSF-A8456 Sigma-) and cells were lysed by lysozyme and sonication. 0.5 ml slurry of HisPur^TM^ Ni-NTA resin equilibrated in lysis buffer was added to the cleared lysate and rotated for 1 h at 4°C before washing with 50 ml lysis buffer and eluted in lysis buffer + 250 mM imidazole. Eluate was concentrated to ∼ 0.4 ml with a 10KDa Amicon Ultra Centrifugal filter and separated in a Superdex200 Increase 10/300 GL column equilibrated in storage buffer (50 mM Tris-HCl pH 8, 0.01% NP40, 10% Glycerol, 500 mM NaCl, 5 mM MgOAc, 0.5 mM TCEP). Peak fractions were pooled, and aliquots were frozen in liquid nitrogen and stored at −80C. Ulp1 eluted at ∼ 13 ml, and protein concentration was determined by Bradford at 2 mg/ml.

### RiboGreen exonuclease assay

For substrate preparation, oligos **i** (unlabelled) and **iii** (unlabelled) (Supplementary Table S4) were diluted to 100 μM in RNase-free water, then mixed to a final concentration of 10 μM each in 20 mM Tris-HCl (pH 7.5) and 50 mM NaCl in 50 μl. Mix was heated at 98°C for 2 min and then cooled to 4°C in 700 cycles of 4 sec each (−0.1°C/cycle).

End-point nuclease assays (20 μl) were performed at room temperature (RT) in reaction buffer 1 (25 mM Tris-HCl pH 7.5, 0.02 % Tween20, 10 % glycerol, 1.5 mM MgCl_2_, 20 mM NaCl, 0.1 mg/ml BSA and 0.5 mM TCEP). Enzyme and substrate were diluted in reaction buffer 1 at the concentration specified in each figure legend. Reactions were stopped by the addition of 30 μl Stop/RiboGreen mix (25 mM Tris-HCl pH 7.5, 0.1 mg/ml BSA, 20 mM EDTA and a 1/250 dilution of Quant-iT™ RiboGreen® RNA reagent, ThermoFisher Scientific) after which fluorescence was read in black 384-well plates (GRE384fb 781076) using a Spark Multimode microplate reader (Tecan) with the following settings: Excitation 485 nm / Emission 530 nm (20/20 bandwith), Gain 94, 30 flashes, Z position of 18843.

### Cy5 gel-based exonuclease assay

Substrate preparation was performed as for the RiboGreen assay, using oligos **ii** (Cy5) and **iii** (unlabelled) (Supplementary Table S4). End-point nuclease assays were performed essentially as the RiboGreen assays except that reactions were stopped (1:1) with 2X sample buffer (98% Formamide, 10 mM EDTA). Samples were denatured at 95°C for 2 min and loaded on 7M-urea polyacrylamide-1X TBE denaturing gels and run in 1x TBE running buffer at 180 V for 2 to 3 hours at RT before Cy5 fluorescence was visualised with an Amersham Imager 600, GE lifesciences.

### Cy3/quencher-based exonuclease assay (kinetic)

For Cy3-substrates preparation, RNA oligos **iv** (Cy3) and **v** (quencher) or DNA oligos **vi** and **vii** were mixed at a 1:1.2 ratio with Cy3 oligo to 20 μM and Iowa Black RQ quencher oligo to 24 μM (Supplementary Table S4). For RNA substrate, samples were heated at 75°C for 3 min and then cooled to 5°C in 700 cycles of 4 seconds each (−0.1°C/cycle) after which substrate was aliquoted and stored at −20°C. For DNA substrate, samples were heated at 95°C for 5 min and then cooled down to 21°C in 740 cycles of 1 second each (−0.1°C/cycle), aliquoted and stored at −20°C.

Cy3/quencher-based kinetic nuclease assays (20 μl) were performed at room temperature (RT) in reaction buffer 2 (50 mM Tris-HCl pH 7.5, 0.01 % Tween20, 10 % glycerol, 1.5 mM MgCl_2_, 20 mM NaCl, 0.1 mg/ml BSA and 0.5 mM TCEP). Enzyme and substrate were diluted in reaction buffer 2 at the concentration specified in each figure legend. Progression of the reactions was monitored every min by fluorescence in black 384-well plates (GRE384fb 781076) using a Spark Multimode microplate reader (Tecan) with the following settings: Excitation 545 nm / Emission 575 nm (10/10 bandwith), Gain 147, 10 flashes, Z position of 17800.

### Methyltransferase assay

The methyltransferase activity of nsp10-14 was assayed by the detection of released SAH from the methyltransferase reaction. Released SAH was detected through the use of the commercially available EPIgeneous™ methyltransferase kit (CisBio Bioassays). Individual kit reagents were reconstituted according to the manufacturer’s instruction. The methyltransferase reaction was conducted at room temperature in an 8 μl reaction volume with 10 nM nsp10-14, 1 μM Ultrapure SAM (CisBio), 0.14 mM GpppA RNA cap analogue (New England Biolabs) in reaction buffer consisting of HEPES-KOH pH 7.6, 150 mM NaCl, and 0.5 mM DTT. The reaction was started with the addition of nsp10-14 and was allowed to proceed for 20 minutes before quenching by the addition of 2 μl 5M NaCl to a final concentration of 1M.

Following quenching, 2 μl Detection Buffer 1 (CisBio) was immediately added to the reaction mixture. After 10 minutes, 4 μl of 16X SAH-d2 conjugate solution (CisBio) was added. 16X SAH-d2 was prepared by adding one part SAH-d2 to 15 parts Detection Buffer 2 (CisBio). After 5 minutes, 4 μl of 1X α-SAH Tb Cryptate antibody solution was added to the reaction mixture. 1X α-SAH Tb Cryptate antibody solution was prepared by adding one part α-SAH Tb Cryptate antibody (CisBio) to 49 parts Detection Buffer 2 (CisBio).

Homogenous Time Resolved Fluorescence (HTRF) measurements were taken after 1 hour following α-SAH Tb Cryptate antibody addition on a Tecan Infinite M1000 Pro plate reader. Readings were taken with a lag time of 60 μs after excitation at λ=337 nm. Readings were taken at emission wavelengths of λ=665nm and λ=620nm. The experimental HTRF ratio (HTRF_exp_) was then calculated as ratio of emission intensities: λ=665/λ=620. To reach the normalised HTRF ratio, HTRF ratio measurements were also taken of wells without enzyme (E_0_) and without SAH-d2 (d2_0_), representing the maximum and minimum achievable HTRF values, respectively. The normalised HTRF ratio was then calculated as a linear transformation of the experimental HTRF ratio, the E_0_ ratio, and the d2_0_ ratio:

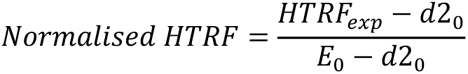

### Screening library

High-throughput screening was performed using a custom compound collection assembled from commercial sources (Sigma, Selleck, Enzo, Tocris, Calbiochem, and Symansis). 2.5 or 7.5 nl of a 10 mM stock of the compounds dissolved in DMSO were arrayed and dispensed into square flat-bottom black 384-well plates containing 1 µl DMSO/well using an Echo 550 (Labcyte), before being sealed and stored at −80°C.

### RiboGreen-based screen (first)

All steps and reagents were kept at room temperature (RT) during the day of the screen. The morning of the screen all plates were moved from −80°C to 4°C. From there, plates were moved to RT 30 min before centrifuging and de-sealing. The RiboGreen screen was performed in three main steps which required the use of three independent XRD-384 Reagent Dispensers (FluidX Ltd.) each loaded with 16-channel tubing and set at high speed. First, 10 μl of 2X enzyme mix (10 nM nsp10-14 fusion protein in 1X reaction buffer 1) was dispensed in columns 1 to 23 and incubated “Enzyme + drugs” for 10 min. After each dispensing, plates were centrifuged for <1 min at 4000 rpm. Second, 10 μl of 2X substrate mix (100 nM unlabelled dsRNA substrate in 1x reaction buffer 1) was dispensed in columns 2 to 24, and 10 μl reaction buffer 1 were pipetted by hand to control columns 1 and 24. After 5 min, 30 μl of Stop/RiboGreen mix (final concentration in 50 µl: 25 mM Tris-HCl pH 7.5, 0.1 mg/ml BSA, 20 mM EDTA and a 4000-fold dilution of Quant-iT™ RiboGreen® RNA reagent) was dispensed in columns 1 to 24. After 2 min, fluorescence in all wells was read using a Spark Multimode microplate reader (Tecan) with the following settings: Excitation 485 nm / Emission 530 nm (20/20 bandwith), Gain 94, 30 flashes, Z position of 18843. Plates were processed one after the other, with ∼ 7 min delay between them.

### Cy3/quencher-based kinetic screen (second)

All steps and reagents were kept at RT during the days of the screen except the stock of enzyme mix that was kept in ice. The morning of the screen, all plates were moved from - 80°C to 4°C. From there, plates were moved to room temperature for 30 min before centrifuging and de-sealing. This second screen was performed in two main steps which required the use of two independent XRD-384 Reagent Dispensers (FluidX Ltd.) each loaded with 16-channel tubing and set at high speed. First, 10 μl of 2X enzyme mix (1 nM nsp10-14 fusion enzyme in 1X reaction buffer 2) was dispensed in columns 1 to 23, and 10 μl reaction buffer 2 were pipetted by hand to control columns 1 and 24. After 10 min, 10 μl of 2X substrate mix (100 nM Cy3/quencher RNA substrate in 1x reaction buffer 2) was dispensed in columns 2 to 24 and, after 2 min, started reading fluorescence in all wells every 1 min for 10 min using a Spark Multimode microplate reader (Tecan) with the following settings: Excitation 545 nm / Emission 575 nm (10/10 bandwith), Gain 147, 10 flashes, Z position of 17800. Plates were processed one after the other, with ∼ 20 min delay between them.

### Screen data analysis

Screen data were analysed with custom MATLAB and R scripts. For the <u>RiboGreen screen</u>, normalised residual activity was calculated for each well, relative to the controls on that plate. The median background fluorescence, calculated from 8 wells containing no substrate, was subtracted from all wells, before normalisation using the formula:

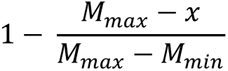

where *M_max_* is the median of 8 wells without enzyme (maximum signal), *M_min_* is the median of 16 wells with no drug (minimum signal) and *x* is the experimental value.

For the <u>second Cy3/quencher-based screen</u>, the slope of each reaction over the first 6 min was calculated by linear regression and then was normalised by dividing by the average of the control wells without drugs in each row of the plate.

In both screens, Z scores were then calculated for each experimental well.

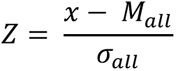

where *x* is the normalised experimental value, and *M_all_* and *σ_all_* are the median and standard deviation of all 6106 normalised samples at that concentration respectively.

Z’ factors were calculated for each plate to determine screen quality.

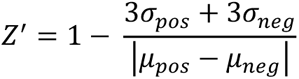

where *μ_pos_* and *σ_pos_* are the mean and standard deviation of the positive controls (no drugs) respectively, and *μ_neg_* and *σ_neg_* are the mean and standard deviation of the negative controls (no enzyme) respectively.

### K_M_ and IC_50_ calculation

For the <u>unlabelled dsRNA substrate</u>, we calculated the initial velocity of the reactions considering 0, 3 and 6 minutes in the presence of 5 nM of nsp10-14 fusion protein and 15.65 - 250 nM substrate by linear regression. Slopes were then used to calculate K_M_ and V_MAX_ by non-linear fitting to the Michaelis-Menten equation using GraphPad Prism.

For the <u>Cy3-dsRNA substrate</u>, we determined the maximum velocity over the first 45 minutes of the reactions in the presence of 0.5 nM of nsp10-14 fusion protein and 15.65 - 500 nM substrate by calculating the maximum of the first derivative. Slopes were then used to calculate K_M_ and V_MAX_ by non-linear fitting to the Michaelis-Menten equation using GraphPad Prism.

Similarly, we determined maximum velocity of the reactions in the presence of 0.5 nM nsp10-14, 50 nM Cy3/Q-substrate and a titration of patulin or ATA (0 - 160 µM) over 15 min by calculating the maximum of the first derivative. Slopes were then used to calculate the percentage of activity relative to the slope without inhibitor. Percent activity for each log_10_ of the concentration of patulin or ATA were then used to estimate the IC_50_ and Hill slopes using GraphPad Prism.

### SARS-CoV-2 production

Batches of the BetaCoV/England/02/2020 (Public Health England) strain of the SARS-CoV-2 virus were produced as in our accompanying manuscripts using VERO E6 cells. A 6-well plate plaque assay was then used to determine plaque forming units (PFU) per ml.

### Viral infectivity assay

Viral infectivity assays were performed as in our accompanying manuscripts. Briefly, 96-well imaging plates (Greiner 655090) were seeded with VERO E6 cells and cultured overnight. The next day, the media was replaced with fresh growth media, followed by addition of drug compounds. Finally, the cells were infected by SARS-CoV2 with a final MOI of 0.5 PFU/cell. 22 h post infection, cells were fixed, permeabilised, and stained for SARS-CoV2 N protein using Alexa488-labelled-CR3009 antibody and cellular DNA using DR (ABCAM). Imaging was carried out using an Opera Phenix (Perkin Elmer) and fluorescent areas and intensity calculated using the Phenix-associated software Harmony (Perkin Elmer). Alexa488/N intensities were normalised to DR/DNA, and to vehicle only samples.

## Data Availability Statement

All data associated with this paper will be deposited in FigShare (https://figshare.com/).

## Author Contributions

**Berta Canal:** Conceptualisation, Methodology, Validation, Formal analysis, Investigation, Resources, Writing – Original Draft, Writing – Review and Editing, Visualisation. **Allison W. McClure:** Conceptualisation, Methodology, Validation, Formal analysis, Investigation, Resources, Writing – Original Draft, Writing – Review and Editing, Visualisation. **Joseph F. Curran:** Conceptualisation, Methodology, Validation, Formal analysis, Investigation, Resources, Writing – Original Draft, Writing – Review and Editing, Visualisation. **Mary Wu:** Methodology, Investigation, Resources. **Rachel Ulferts:** Methodology, Investigation. **Florian Weissmann:** Resources. **Jingkun Zeng:** Resources, Software. **Agustina P. Bertolin:** Resources. **Jennifer C. Milligan:** Investigation, Software. **Souradeep Basu:** Investigation. **Lucy S. Drury:** Investigation. **Tom Deegan:** Resources. **Ryo Fujisawa:** Resources. **Emma L. Roberts:** Resources. **Clovis Basier:** Resources. **Karim Labib:** Supervision. **Rupert Beale:** Supervision. **Michael Howell:** Supervision**. John F.X Diffley:** Conceptualisation, Methodology, Writing – Review and Editing, Supervision, Project administration, Funding acquisition.

## Acknowledgements

We thank the Crick high-throughput screen (HTS) science technology platform (STP) for providing the chemical library and for help with screen design and analysis and are grateful to MRC Reagents and Services (https://mrcppureagents.dundee.ac.uk/) for providing DU70487 DNA construct for nsp14. We thank the Fermentation unit of the Crick Structural biology STP for large scale protein expression. We thank David McClure for help with screen analysis. We thank Anne Early for ordering supplies and hit compounds. We thank Chris Smith for help with initial screen design and concepts. This work was supported by the Francis Crick Institute, which receives its core funding from Cancer Research UK (FC001066), the UK Medical Research Council (FC001066), and the Wellcome Trust (FC001066). This work was also funded by a Wellcome Trust Senior Investigator Award (106252/Z/14/Z) to J.F.X.D. BC and FW have received funding from the European Union’s Horizon 2020 research and innovation programme under the Marie Skłodowska-Curie grant agreement Nos 895786 and 844211. JZ has received funding from a Ph.D. fellowship awarded by Boehringer Ingelheim Fonds.

## Supplementary Material

**Supplementary Figure S1.**
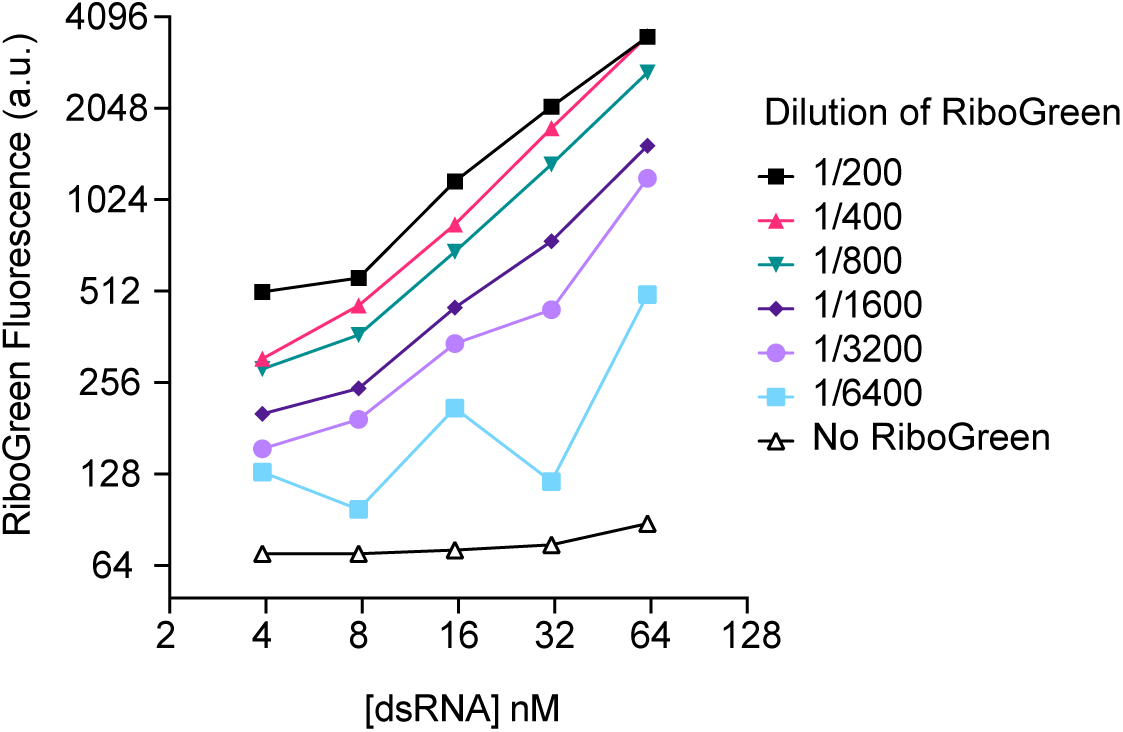
Titration of RiboGreen. RiboGreen was diluted as indicated and then incubated with 4 to 64 nM unlabelled dsRNA substrate prior to fluorescence detection.

**Supplementary Figure S2.**
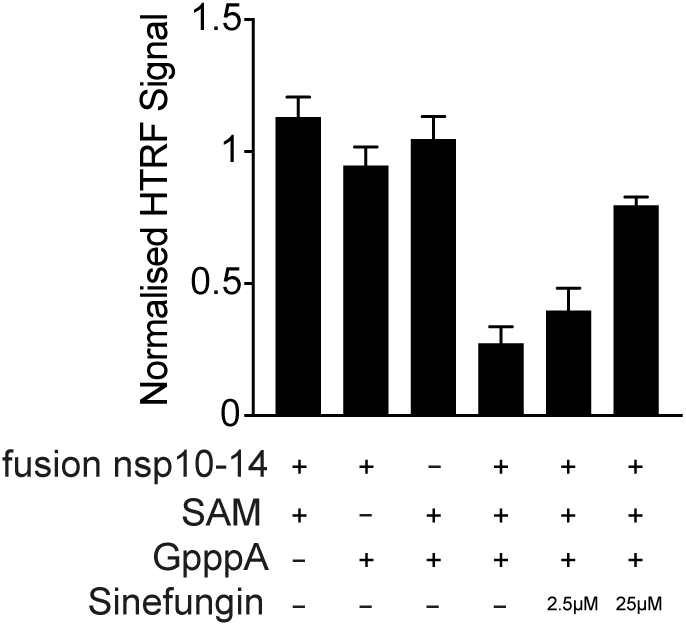
The nsp10-14 fusion functions as a methyltransferase. The nsp10-14 fusion protein was assayed for methyltransferase activity by the detection of formed SAH following methyltransferase assay (see Experimental Procedures). The methyltransferase reaction was run in either the absence of 10 nM nsp10-14, 1 μM SAM methyl donor, 0.11 mM GpppA cap analogue, or in the presence of all three components. In addition, the methyltransferase reaction was conducted in the presence of 2.5 μM and 25 μM of the pan-methyltransferase inhibitor Sinefungin, which acts as a competitive inhibitor (with respect to SAM) towards SAM-dependent methyltransferases. Error bars represent standard deviation from the mean.

**Supplementary Figure S3.**
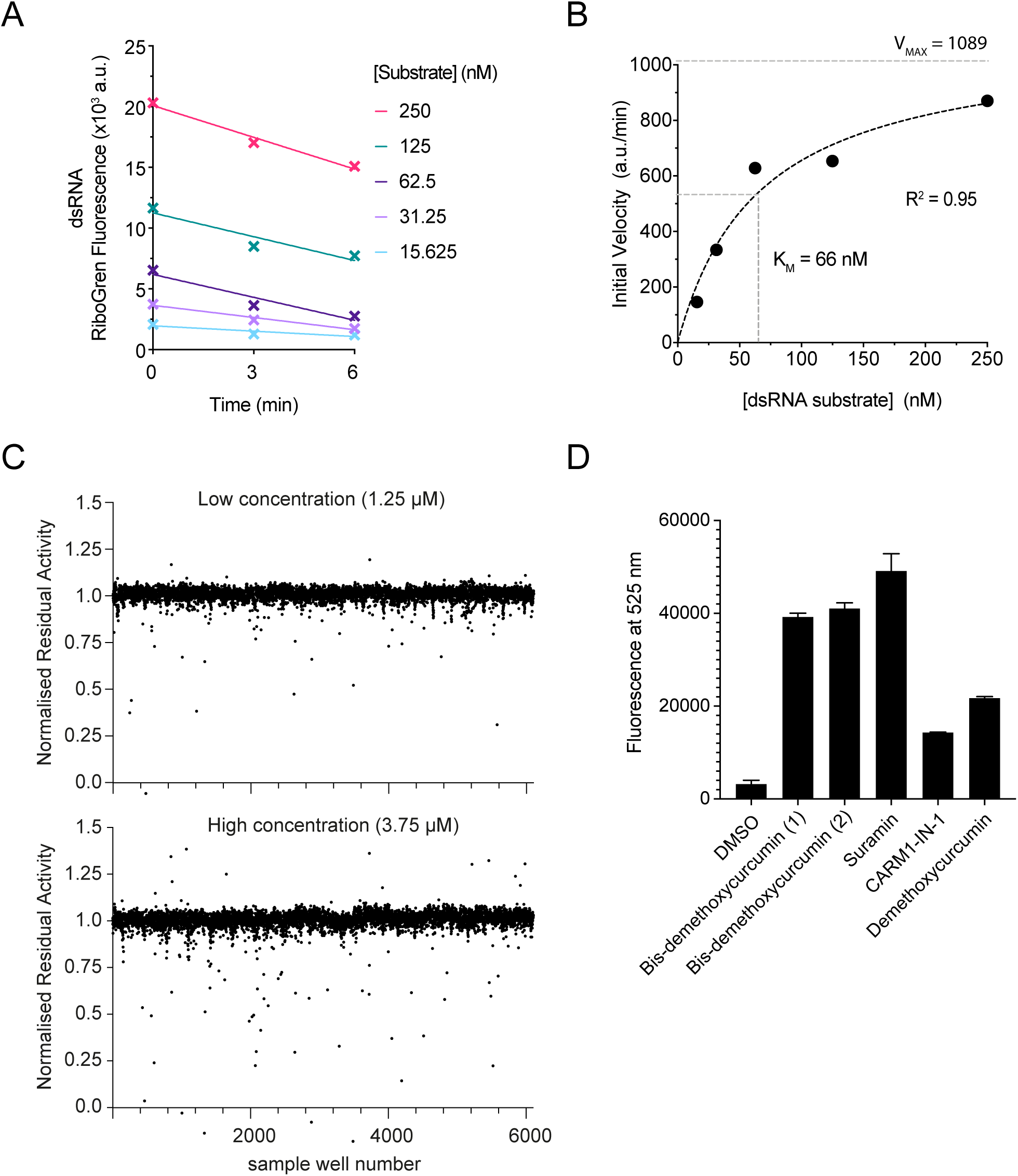
Ribogreen-based screen to identify SARS-CoV-2 nsp14/10 exoribonuclease inhibitors. **A.** Determination of initial rate of reaction when varying unlabelled dsRNA substrate concentration (15 – 250 nM). Reactions were carried out using 5 nM nsp10-14 fusion enzyme and stopped after 0, 3 and 6 minutes before dsRNA detection with RiboGreen. **B.** Non-linear Michaelis-Menten fit for enzyme kinetics data generated by substrate titration from A. **C.** Normalised residual activity for screen samples at low and high concentrations. **D.** Selected compounds were assayed for autofluorescence at 525 nm (RiboGreen wavelength). Error bars represent standard deviation from the mean.

**Supplementary Figure S4.**
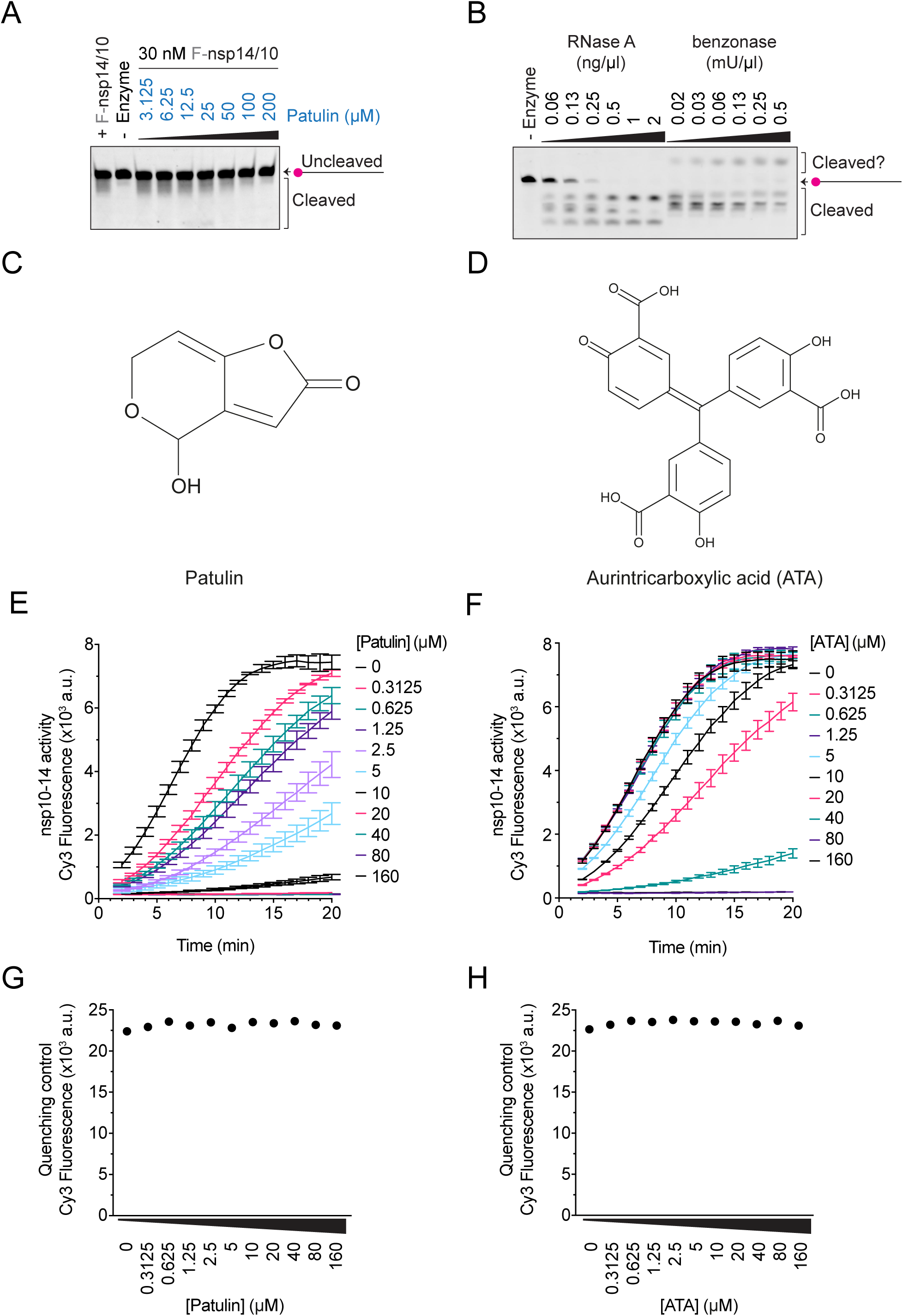
Patulin and aurintricarboxylic acid inhibit nsp14/10 nuclease in vitro. **A.** 30 nM of co-expressed nsp14/10 complex was pre-incubated with 3 - 200 µM of patulin, and nuclease reactions were performed in the presence of 50 nM Cy5-dsRNA substrate at RT for 5 min and visualised in TBE-urea polyacrylamide gels. **B.** Nuclease reactions containing titration of RNase A or benzonase and 50 nM Cy5-dsRNA substrate were performed at RT for 5 minutes and visualised by TBE-urea polyacrylamide gels. **C, D.** Representation of the chemical structures of patulin (C) and ATA (D). **E, F.** Kinetic nuclease reactions (20 min) were performed in triplicates in the presence of 0.5 nM nsp10-14 fusion, 50 nM Cy3-dsRNA substrate and 0 to 160 µM patulin (C) or ATA (D). Patulin and ATA were pre-incubated with nsp10-14 for 10 minutes prior to the addition of the substrate. These reactions were used to calculate the IC_50_ of the inhibitors (see Experimental Procedures). Error bars represent standard deviation from the mean. **G, H.** Drug quenching test performed by assessing fluorescence of a Cy3-labelled oligonucleotide (50 nM) pre-incubated 10 minutes in the presence of a titration (0 – 160 µM) of patulin and ATA respectively.

**Supplementary Figure S5.**
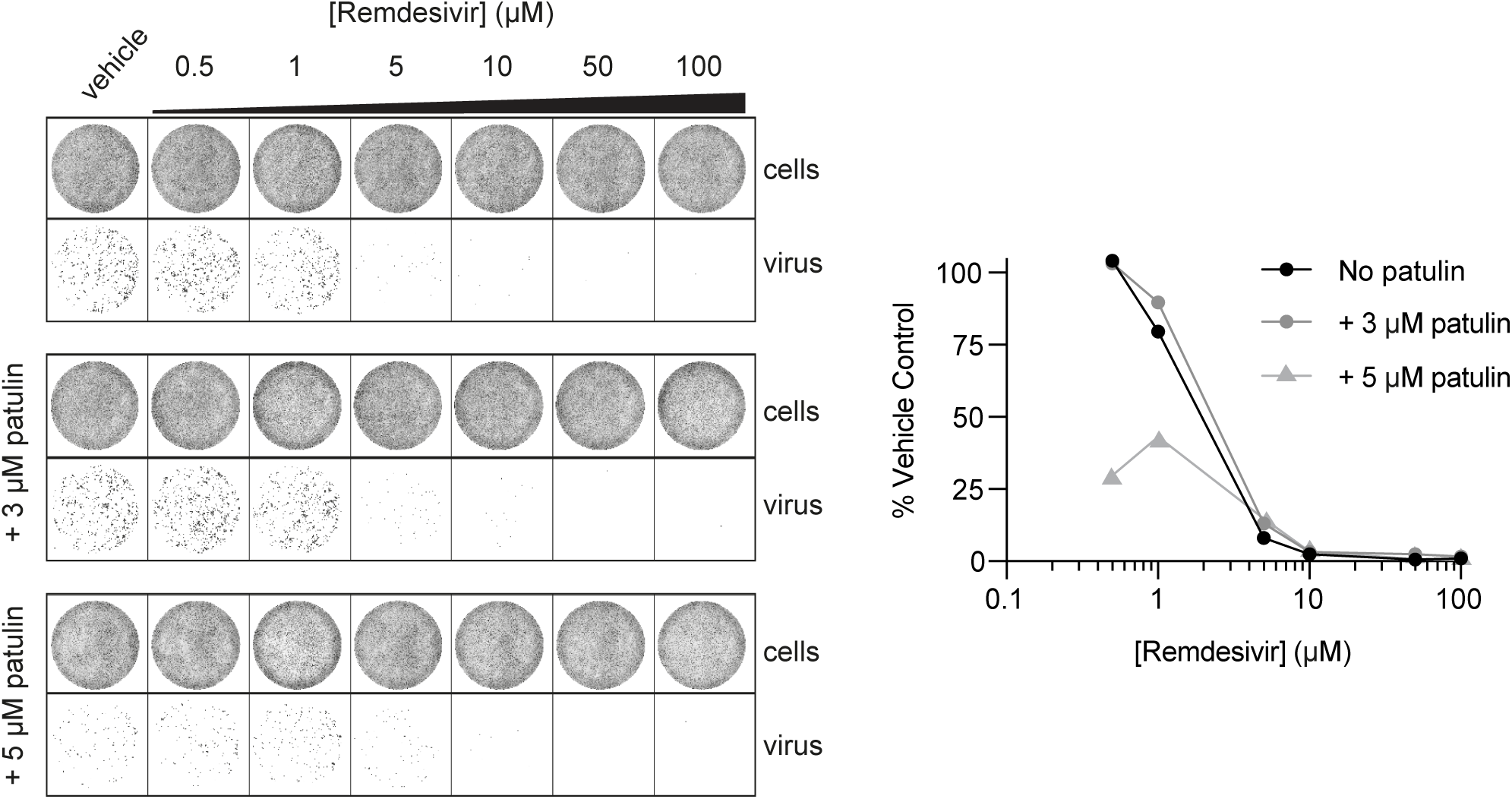
Patulin and remdesivir are not synergistic to inhibit SARS-CoV-2 viral growth in VERO cells. SARS-CoV-2 infectivity assays in VERO E6 assay performed as in Figure 5A with the indicated concentrations of patulin and remdesivir. For each condition, quantifications represent viral area normalised to the no drug well. Error bars represent standard deviation from the mean.

**Supplementary Table S1.**
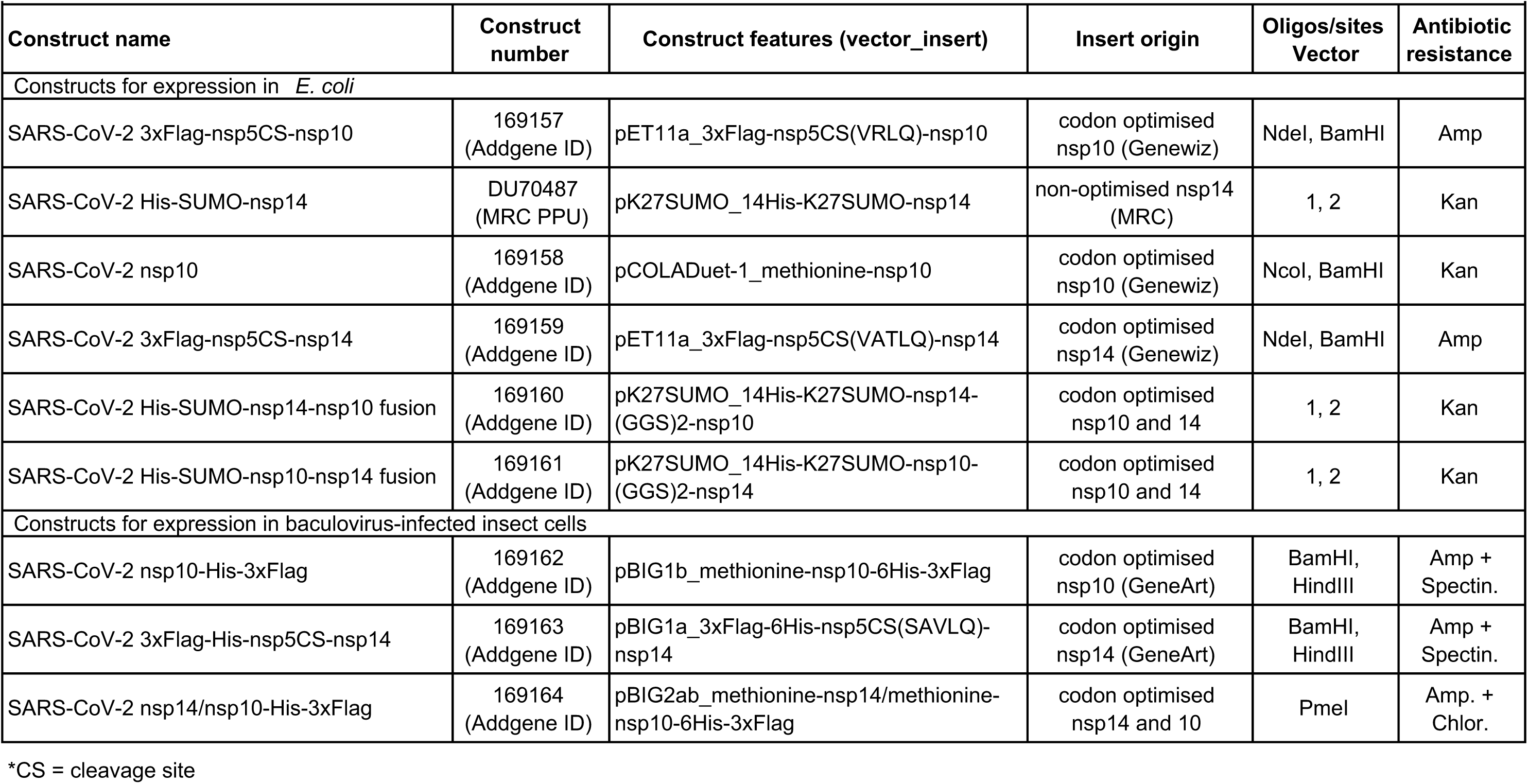
Cloning strategies

**Supplementary Table S2.**
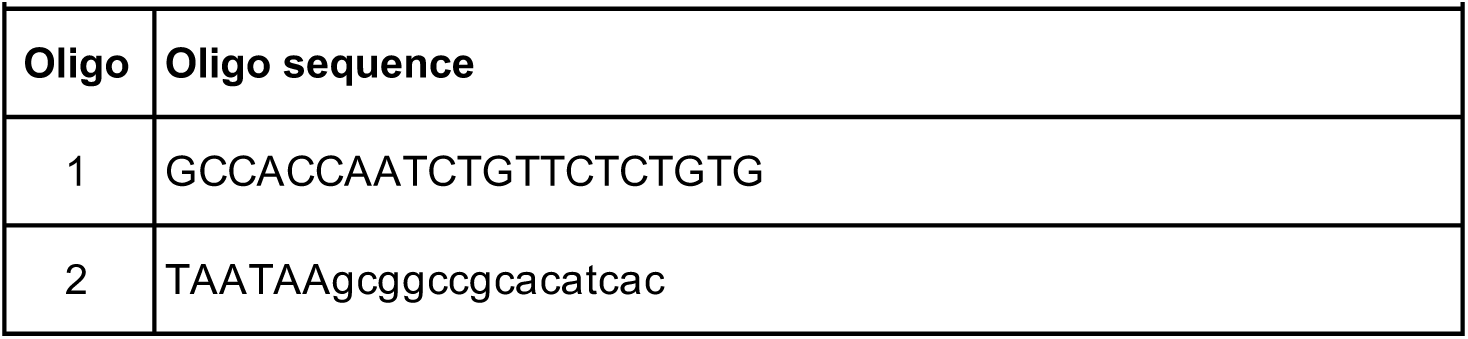
Cloning oligos

**Supplementary Table S3.** Purification strategies

| Supplementary Table S3. Purification strategies |  |  |  |
| --- | --- | --- | --- |
| Plasmid name | Used to express | Codon optimisation | Purification steps |
| SARS-CoV-2 3xFlag-nsp5CS-nsp10 | Individual nsp10 | *E. coli* | PD α-Flag > MonoQ > Dialysis |
| SARS-CoV-2 His-SUMO-nsp14 | Individual nsp14 | Non-optimised | PD NiNTA > Ulp1 > Superdex200 |
| SARS-CoV-2 nsp10 | Complex nsp14/10 | *E. coli* | PD α-Flag > MonoQ > Superdex200 |
| SARS-CoV-2 3xFlag-nsp5CS-nsp14 |  | *E. coli* |  |
| SARS-CoV-2 His-SUMO-nsp14-nsp10 fusion | nsp14-10 fusion | *E. coli* | PD NiNTA > Ulp1 > MonoQ > Superdex200 |
| SARS-CoV-2 His-SUMO-nsp10-nsp14 fusion | nsp10-14 fusion | *E. coli* | PD NiNTA > Ulp1 > MonoQ > Superdex200 |
| SARS-CoV-2 nsp10-His-3xFlag | Individual nsp10 | Insect cells | PD α-Flag > MonoQ > Dialysis |
| SARS-CoV-2 3xFlag-His-nsp5CS-nsp14 | Individual nsp14 | Insect cells | PD α-Flag > Superdex200 |
| SARS-CoV-2 nsp14/nsp10-His-3xFlag | Complex nsp14/10 | Insect cells | PD α-Flag > MonoQ > Superdex200 |

**Supplementary Table S4.**
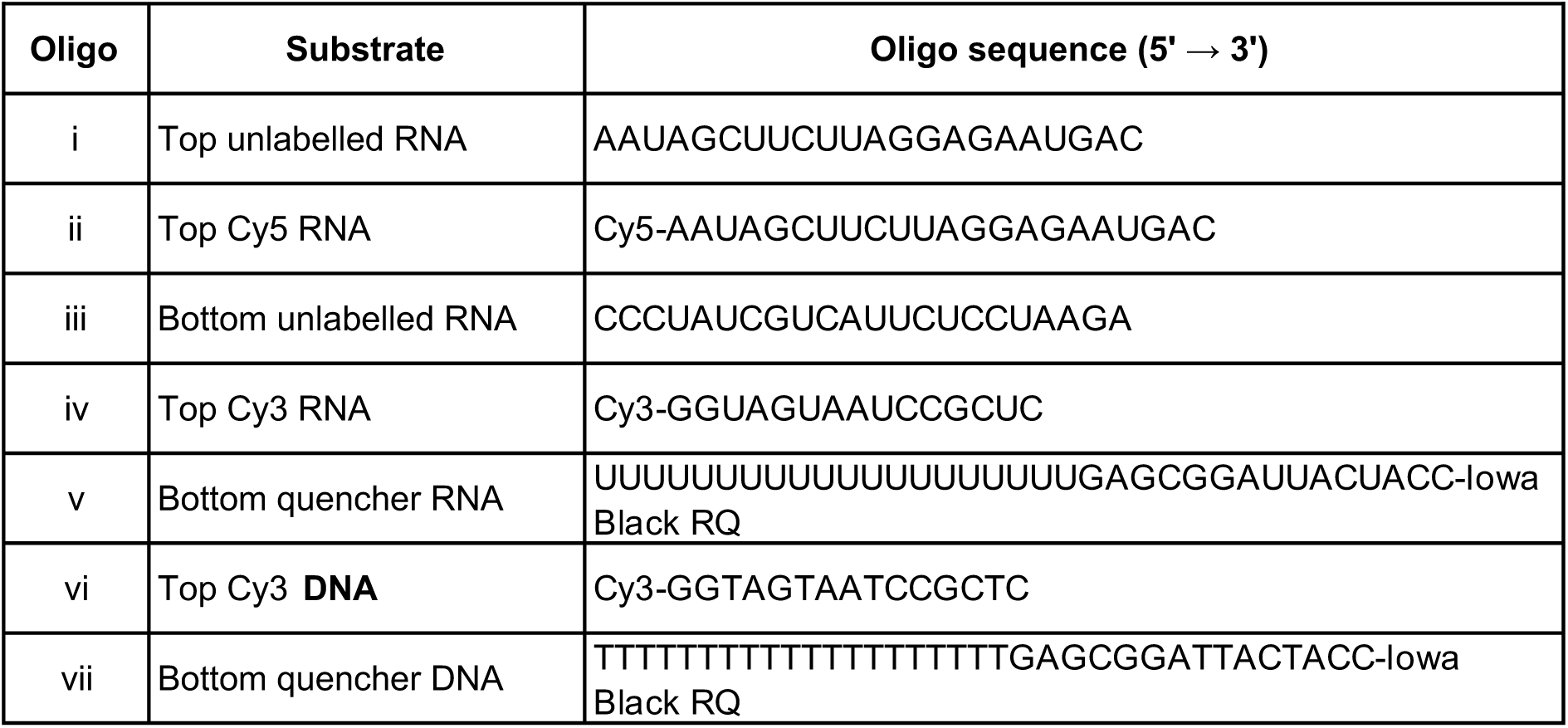
Substrate oligos

**Supplementary Table S5.**
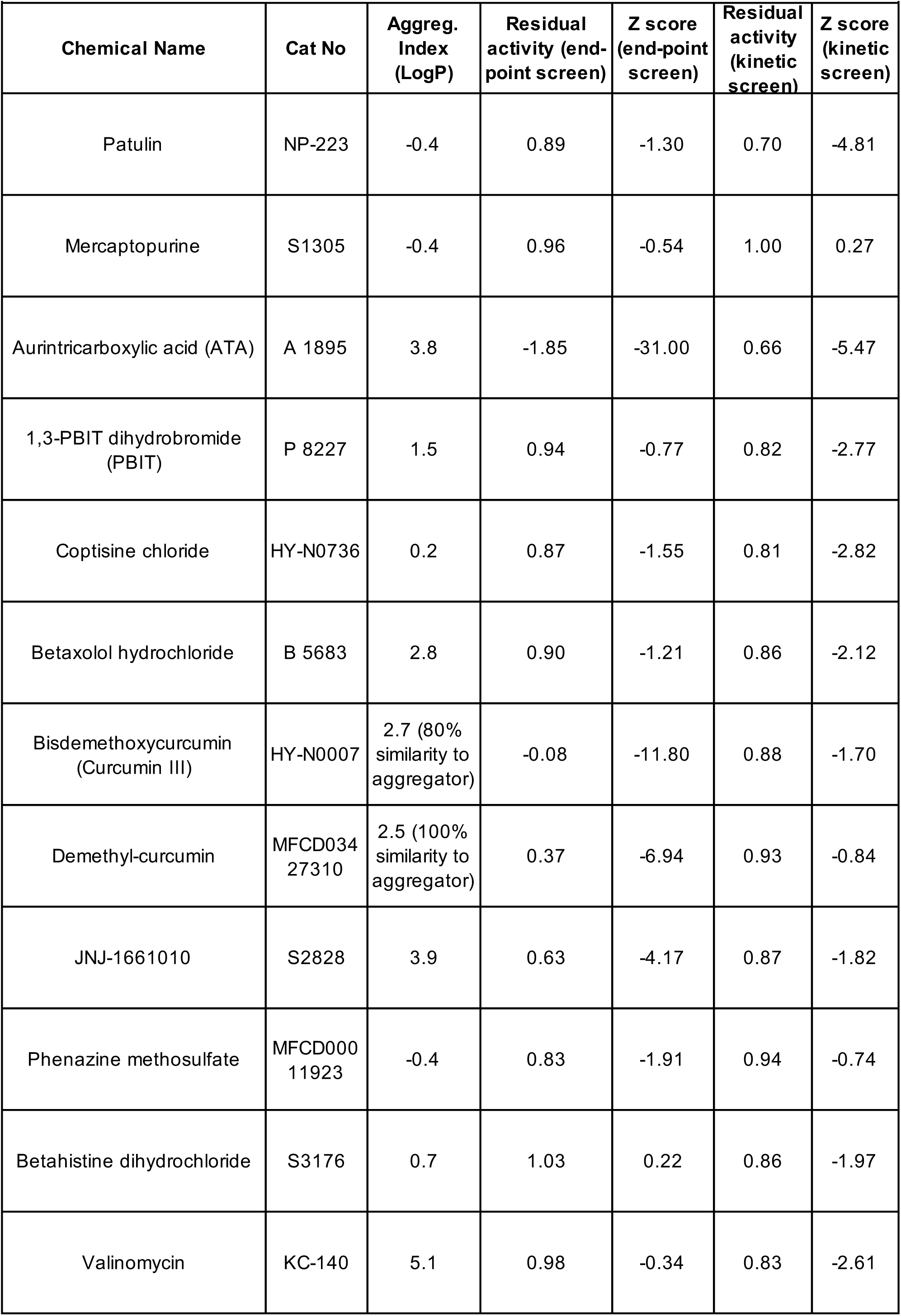
Top screen hit compounds

## References

1. The species Severe acute respiratory syndrome-related coronavirus: classifying 2019-nCoV and naming it SARS-CoV-2. Nat Microbiol, 2020. 5(4): p. 536–544.

2. Zhou, P., et al., A pneumonia outbreak associated with a new coronavirus of probable bat origin. Nature, 2020. 579(7798): p. 270–273.

3. Xiong, T.Y., et al., Coronaviruses and the cardiovascular system: acute and long-term implications. Eur Heart J, 2020. 41(19): p. 1798–1800.

4. Zhao, Y.M., et al., Follow-up study of the pulmonary function and related physiological characteristics of COVID-19 survivors three months after recovery. EClinicalMedicine, 2020. 25: p. 100463.

5. Huerga Encabo, H., et al., Human Erythroid Progenitors Are Directly Infected by SARS-CoV-2: Implications for Emerging Erythropoiesis in Severe COVID-19 Patients. Stem Cell Reports, 2021. 16(3): p. 428–436.

6. Hu, B., et al., Characteristics of SARS-CoV-2 and COVID-19. Nature Reviews Microbiology, 2021. 19(3): p. 141–154.

7. Totura, A.L. and S. Bavari, Broad-spectrum coronavirus antiviral drug discovery. Expert Opin Drug Discov, 2019. 14(4): p. 397–412.

8. Medline, A., et al., Evaluating the impact of stay-at-home orders on the time to reach the peak burden of Covid-19 cases and deaths: does timing matter? BMC Public Health, 2020. 20(1): p. 1750.

9. Cui, J., F. Li, and Z.-L. Shi, Origin and evolution of pathogenic coronaviruses. Nature Reviews Microbiology, 2019. 17(3): p. 181–192.

10. Peiris, J.S., et al., Coronavirus as a possible cause of severe acute respiratory syndrome. Lancet, 2003. 361(9366): p. 1319–25.

11. Zaki, A.M., et al., Isolation of a novel coronavirus from a man with pneumonia in Saudi Arabia. N Engl J Med, 2012. 367(19): p. 1814–20.

12. Agostini, M.L., et al., Coronavirus Susceptibility to the Antiviral Remdesivir (GS-5734) Is Mediated by the Viral Polymerase and the Proofreading Exoribonuclease. mBio, 2018. 9(2).

13. Wang, M., et al., Remdesivir and chloroquine effectively inhibit the recently emerged novel coronavirus (2019-nCoV) in vitro. Cell Res, 2020. 30(3): p. 269–271.

14. Yin, W., et al., Structural basis for inhibition of the RNA-dependent RNA polymerase from SARS-CoV-2 by remdesivir. Science, 2020. 368(6498): p. 1499–1504.

15. Beigel, J.H., et al., Remdesivir for the Treatment of Covid-19 - Final Report. N Engl J Med, 2020. 383(19): p. 1813–1826.

16. Pan, H., et al., Repurposed Antiviral Drugs for Covid-19 - Interim WHO Solidarity Trial Results. N Engl J Med, 2020.

17. V’Kovski, P., et al., Coronavirus biology and replication: implications for SARS-CoV-2. Nat Rev Microbiol, 2020: p. 1–16.

18. Finkel, Y., et al., The coding capacity of SARS-CoV-2. Nature, 2021. 589(7840): p. 125–130.

19. Sevajol, M., et al., Insights into RNA synthesis, capping, and proofreading mechanisms of SARS-coronavirus. Virus Res, 2014. 194: p. 90–9.

20. Minskaia, E., et al., Discovery of an RNA virus 3’->5’ exoribonuclease that is critically involved in coronavirus RNA synthesis. Proc Natl Acad Sci U S A, 2006. 103(13): p. 5108–13.

21. Chen, Y., et al., Functional screen reveals SARS coronavirus nonstructural protein nsp14 as a novel cap N7 methyltransferase. Proc Natl Acad Sci U S A, 2009. 106(9): p. 3484–9.

22. Bouvet, M., et al., In vitro reconstitution of SARS-coronavirus mRNA cap methylation. PLoS Pathog, 2010. 6(4): p. e1000863.

23. Ma, Y., et al., Structural basis and functional analysis of the SARS coronavirus nsp14-nsp10 complex. Proc Natl Acad Sci U S A, 2015. 112(30): p. 9436–41.

24. Moser, M.J., et al., The proofreading domain of Escherichia coli DNA polymerase I and other DNA and/or RNA exonuclease domains. Nucleic Acids Res, 1997. 25(24): p. 5110–8.

25. Zuo, Y. and M.P. Deutscher, Exoribonuclease superfamilies: structural analysis and phylogenetic distribution. Nucleic Acids Res, 2001. 29(5): p. 1017–26.

26. Bouvet, M., et al., RNA 3’-end mismatch excision by the severe acute respiratory syndrome coronavirus nonstructural protein nsp10/nsp14 exoribonuclease complex. Proc Natl Acad Sci U S A, 2012. 109(24): p. 9372–7.

27. Gorbalenya, A.E., et al., Nidovirales: evolving the largest RNA virus genome. Virus Res, 2006. 117(1): p. 17–37.

28. Eckerle, L.D., et al., High fidelity of murine hepatitis virus replication is decreased in nsp14 exoribonuclease mutants. J Virol, 2007. 81(22): p. 12135–44.

29. Smith, E.C., et al., Coronaviruses lacking exoribonuclease activity are susceptible to lethal mutagenesis: evidence for proofreading and potential therapeutics. PLoS Pathog, 2013. 9(8): p. e1003565.

30. Graepel, K.W., et al., Proofreading-Deficient Coronaviruses Adapt for Increased Fitness over Long-Term Passage without Reversion of Exoribonuclease-Inactivating Mutations. mBio, 2017. 8(6).

31. Ogando, N.S., et al., The Curious Case of the Nidovirus Exoribonuclease: Its Role in RNA Synthesis and Replication Fidelity. Front Microbiol, 2019. 10: p. 1813.

32. Smith, E.C., et al., Mutations in coronavirus nonstructural protein 10 decrease virus replication fidelity. J Virol, 2015. 89(12): p. 6418–26.

33. Eckerle, L.D., et al., Infidelity of SARS-CoV Nsp14-Exonuclease Mutant Virus Replication Is Revealed by Complete Genome Sequencing. PLOS Pathogens, 2010. 6(5): p. e1000896.

34. Subissi, L., et al., One severe acute respiratory syndrome coronavirus protein complex integrates processive RNA polymerase and exonuclease activities. Proc Natl Acad Sci U S A, 2014. 111(37): p. E3900–9.

35. Yan, L., et al., Cryo-EM Structure of an Extended SARS-CoV-2 Replication and Transcription Complex Reveals an Intermediate State in Cap Synthesis. Cell, 2021. 184(1): p. 184–193.e10.

36. Ogando, N.S., et al., The Enzymatic Activity of the nsp14 Exoribonuclease Is Critical for Replication of MERS-CoV and SARS-CoV-2. J Virol, 2020. 94(23).

37. Becares, M., et al., Mutagenesis of Coronavirus nsp14 Reveals Its Potential Role in Modulation of the Innate Immune Response. J Virol, 2016. 90(11): p. 5399–5414.

38. Case, J.B., et al., Murine Hepatitis Virus nsp14 Exoribonuclease Activity Is Required for Resistance to Innate Immunity. Journal of Virology, 2018. 92(1): p. e01531–17.

39. Gribble, J., et al., The coronavirus proofreading exoribonuclease mediates extensive viral recombination. PLoS Pathog, 2021. 17(1): p. e1009226.

40. Ashburn, T.T. and K.B. Thor, Drug repositioning: identifying and developing new uses for existing drugs. Nature Reviews Drug Discovery, 2004. 3(8): p. 673–683.

41. Guy, R.K., et al., Rapid repurposing of drugs for COVID-19. Science, 2020. 368(6493): p. 829–830.

42. Irwin, J.J., et al., An Aggregation Advisor for Ligand Discovery. J Med Chem, 2015. 58(17): p. 7076–87.

43. Richard B. Hallick, B.K.C., Patrick W. Gray and Emil M. Orozco, Jr, Use of aurintricarboxylic acid as an inhibitor of nucleases during nucleic acid isolation. 1977.

44. Zheng, W., W. Sun, and A. Simeonov, Drug repurposing screens and synergistic drug-combinations for infectious diseases. British Journal of Pharmacology, 2018. 175(2): p. 181–191.

45. Bouvet, M., et al., Coronavirus Nsp10, a critical co-factor for activation of multiple replicative enzymes. J Biol Chem, 2014. 289(37): p. 25783–96.

46. Saramago, M., et al., New targets for drug design: Importance of nsp14/nsp10 complex formation for the 3’-5’ exoribonucleolytic activity on SARS-CoV-2. bioRxiv, 2021: p. 2021.01.07.425745.

47. Totura, A.L. and R.S. Baric, SARS coronavirus pathogenesis: host innate immune responses and viral antagonism of interferon. Current Opinion in Virology, 2012. 2(3): p. 264–275.

48. Zhang, Q., K. Shi, and D. Yoo, Suppression of type I interferon production by porcine epidemic diarrhea virus and degradation of CREB-binding protein by nsp1. Virology, 2016. 489: p. 252–68.

49. Robson, F., et al., Coronavirus RNA Proofreading: Molecular Basis and Therapeutic Targeting. Mol Cell, 2020. 79(5): p. 710–727.

50. Gonzalez, R.G., R.S. Haxo, and T. Schleich, Mechanism of action of polymeric aurintricarboxylic acid, a potent inhibitor of protein-nucleic acid interactions. Biochemistry, 1980. 19(18): p. 4299–4303.

51. Baddock, H.T., et al., Characterisation of the SARS-CoV-2 ExoN (nsp14-ExoN-nsp10) complex: implications for its role in viral genome stability and inhibitor identification. bioRxiv, 2020: p. 2020.08.13.248211.

52. He, R., et al., Potent and selective inhibition of SARS coronavirus replication by aurintricarboxylic acid. Biochem Biophys Res Commun, 2004. 320(4): p. 1199–203.

53. Park, J.G., et al., Potent Inhibition of Zika Virus Replication by Aurintricarboxylic Acid. Front Microbiol, 2019. 10: p. 718.

54. Lee, M., et al., Selective inhibition of the membrane attack complex of complement by low molecular weight components of the aurin tricarboxylic acid synthetic complex. Neurobiol Aging, 2012. 33(10): p. 2237–46.

55. Klein, P., et al., In vitro and in vivo activity of aurintricarboxylic acid preparations against Cryptosporidium parvum. Journal of Antimicrobial Chemotherapy, 2008. 62(5): p. 1101–1104.

56. Puel, O., P. Galtier, and I.P. Oswald, Biosynthesis and toxicological effects of patulin. Toxins (Basel), 2010. 2(4): p. 613–31.

57. Arzu Kockaya, E., et al., Evaluation of patulin toxicity in the thymus of growing male rats. Arh Hig Rada Toksikol, 2009. 60(4): p. 411–8.

58. Saxena, N., et al., Role of mitogen activated protein kinases in skin tumorigenicity of patulin. Toxicol Appl Pharmacol, 2011. 257(2): p. 264–71.

59. Saxena, N., et al., Patulin causes DNA damage leading to cell cycle arrest and apoptosis through modulation of Bax, p(53) and p(21/WAF1) proteins in skin of mice. Toxicol Appl Pharmacol, 2009. 234(2): p. 192–201.

60. Seigle-Murandi, F., et al., Antitumor activity of patulin and structural analogs. Pharmazie, 1992. 47(4): p. 288–91.

61. Clarke, M., The 1944 patulin trial of the British Medical Research Council. J R Soc Med, 2006. 99(9): p. 478–80.

62. Stein, A., et al., Key steps in ERAD of luminal ER proteins reconstituted with purified components. Cell, 2014. 158(6): p. 1375–1388.

63. Weissmann, F., et al., biGBac enables rapid gene assembly for the expression of large multisubunit protein complexes. Proc Natl Acad Sci U S A, 2016. 113(19): p. E2564–9.

64. Trowitzsch, S., et al., New baculovirus expression tools for recombinant protein complex production. J Struct Biol, 2010. 172(1): p. 45–54.

